# Patient-Derived Xenograft-Guided Prediction of Chemotherapy Response in Triple-Negative Breast Cancer

**DOI:** 10.1101/2024.12.09.627518

**Authors:** Jonathan T. Lei, Lacey E. Dobrolecki, Chen Huang, Ramakrishnan R. Srinivasan, Suhas V. Vasaikar, Alaina N. Lewis, Christina Sallas, Na Zhao, Jin Cao, John D. Landua, Jacob B. Pilcher, Chang In Moon, Yuxing Liao, Susan G. Hilsenbeck, Mothaffar F. Rimawi, Matthew J. Ellis, Varduhi Petrosyan, Bo Wen, Kai Li, Alexander B. Saltzman, Antrix Jain, Anna Malovannaya, Gerburg M. Wulf, Elisabetta Marangoni, Shunqiang Li, Daniel C. Kraushaar, Tao Wang, Matthew P. Goetz, Judy C. Boughey, Liewei Wang, Krishna R. Kalari, Vera J. Suman, Senthil Damodaran, Xiaofeng Zheng, Funda Meric-Bernstam, Gloria V. Echeverria, Meenakshi Anurag, Xi Chen, Bryan E. Welm, Alana L. Welm, Bing Zhang, Michael T. Lewis

**Affiliations:** Lester and Sue Smith Breast Center and Dan L Duncan Comprehensive Cancer Center, Baylor College of Medicine, Houston, TX 77030, USA; Department of Molecular and Cellular Biology, Baylor College of Medicine, Houston, TX 77030, USA; Department of Molecular and Human Genetics, Baylor College of Medicine, Houston, TX 77030, USA; Mass Spectrometry Proteomics Core, Baylor College of Medicine, Houston, TX 77030, USA; Department of Biochemistry and Molecular Pharmacology, Baylor College of Medicine, Houston, TX 77030, USA; Cancer Research Institute, Department of Medicine, Beth Israel Deaconess Medical Center, Harvard Medical School, Boston, MA 02215, USA; Laboratory of Preclinical Investigation, Translational Research Department, Institut Curie, PSL University, 26 Rue d’Ulm, Paris 75005, France; Siteman Cancer Center, Department of Medicine, Washington University in St. Louis, St. Louis, MO 63108, USA; Department of Oncology, Mayo Clinic, Rochester, MN 55905, USA; Division of Breast and Melanoma Surgical Oncology, Department of Surgery, Mayo Clinic, Rochester, MN, USA; Department of Molecular Pharmacology and Experimental Therapeutics, Mayo Clinic, Rochester, MN 55905, USA; Department of Quantitative Health Sciences, Mayo Clinic, Rochester, MN 55905, USA; MD Anderson Cancer Center, Houston, TX 77030, USA; Department of Medicine, Baylor College of Medicine, Houston, TX 77030, USA; Department of Surgery, Huntsman Cancer Institute, University of Utah, Salt Lake City, UT 84112, USA; Department of Oncological Sciences, Huntsman Cancer Institute, University of Utah, Salt Lake City, UT 84112, USA; Department of Radiology, Baylor College of Medicine, Houston, TX 77030, USA; Department of Genetics, University of Alabama at Birmingham, Birmingham, AL 35294, USA; Translational Oncology Bioinformatics, Pfizer, Bothell, WA 98021, USA; The MOE Key Laboratory of Biosystems Homeostasis & Protection and Innovation Center for Cell Signaling Network, Life Sciences Institute, Zhejiang University, Hangzhou 310058, China; Sage Bionetworks, Seattle, WA, 98121; Research Informatics, Information Technology, Boston Children’s Hospital, Boston, MA 02115, USA; State University of Campinas (UNICAMP), Campinas, São Paulo, Brazil; Department of Genome Sciences, University of Washington, Seattle, WA 98195, USA; Department of Computational Medicine and Bioinformatics, University of Michigan, Ann Arbor, MI 48109, USA

## Abstract

Chemotherapy regimens for triple-negative breast cancer (TNBC) combine agents without knowing which agents drive response. Consequently, predictors derived from multi-agent regimens cannot be assumed to generalize to individual drugs or other regimens, motivating development of treatment-matched predictors. Here, we used TNBC patient-derived xenografts (PDXs) treated with carboplatin, docetaxel, or the combination to deconvolute drug-specific responses and identify associated molecular features. Combination treatment rarely improved upon the best single agent, with enhanced responses in only 13% of PDXs and antagonism in a comparable fraction. Proteogenomic analyses identified high cytokeratin-5 (KRT5) as a general marker of chemotherapy responsiveness and KRT5 immunohistochemistry discriminated responsive PDXs (AUROC, 0.83). To train treatment-specific predictors, we integrated these data with independent PDX and clinical cohorts with responses assessed after anthracycline-free platinum, taxane, or platinum-taxane therapy, ensuring response corresponded to the modeled treatment. Four feature selection strategies yielded largely nonoverlapping biomarker panels converging on treatment-relevant pathways. On independent test data, RNA-based predictors of complete response (CR) to platinum-based (carboplatin or cisplatin) and taxane-based (docetaxel or paclitaxel) chemotherapy achieved AUROCs of 0.80 and 0.86, respectively. For platinum-taxane regimens (carboplatin plus docetaxel or paclitaxel), proteomic-guided feature selection generated a 10-biomarker, protein-informed RNA predictor of pathologic complete response (pCR) that outperformed RNA-only feature selection and achieved an AUROC of 0.85 in an independent clinical cohort, while retaining practicality as an RNA-based assay. Treatment-matched integration of multi-omic PDX and clinical datasets provides a framework for chemotherapy-specific predictors with clinically relevant performance, supporting biomarker-guided precision selection and treatment optimization for patients with TNBC.

**Statement of significance:** Integration of multi-omic data from patient-derived xenografts with treatment-matched clinical cohorts yielded three retrospectively validated predictors of platinum, taxane, and platinum+taxane response, enabling biomarker-guided chemotherapy selection for patients with triple-negative breast cancer.

## Introduction

Breast cancer is a leading cause of cancer-related deaths worldwide. Triple-negative breast cancer (TNBC) is defined by the absence of estrogen receptor (ER) and progesterone receptor (PR) and low/no expression of HER2, which precludes endocrine and HER2-directed therapy. Beyond chemotherapy, two systemic options are now established for patient subsets: 1) immune checkpoint blockade (e.g., pembrolizumab) is added to neoadjuvant chemotherapy for high-risk disease (tumors ≥2 cm or node-positive), where it improves pathologic complete response and event-free survival and 2) PARP inhibition is approved as adjuvant therapy for the ∼15% of patients with TNBC harboring deleterious BRCA1/2 mutations^1^. In both contexts, combination chemotherapy remains the backbone of systemic treatment. In the neoadjuvant setting it consists of taxanes (paclitaxel, docetaxel), anthracyclines (doxorubicin, epirubicin), and cyclophosphamide, often with a platinum (cisplatin, carboplatin), given in series and/or in combination. Identifying patients who will respond to chemotherapy, and defining optimal regimens for TNBC, remain areas of intense investigation.

The addition of taxanes to anthracycline/cyclophosphamide (AC)-based regimens improved outcomes^2,3^, and was the standard-of-care to treat breast tumors for many years. The discovery that a subset of TNBC may be sensitive to DNA damaging agents^4^ prompted the investigation of platinum derivatives for TNBC. More recently, anthracycline-containing neoadjuvant regimens incorporating platinum-taxane combinations have shown promising activity, improving pathologic complete response rates compared with taxane-containing regimens without platinum. For example, although the chemotherapy backbones differed among clinical trials such as GeparSixto^5^, BrighTNess^6^, and CALGB 40603^7^, TNBC patients enrolled in treatment arms receiving platinum- taxane combinations achieved pCR rates exceeding 50%, significantly higher than those in the corresponding control arms.

Because anthracycline-based regimens can increase the frequency of adverse events^8^, more current studies such as NeoCART^9^, and two TNBC clinical trials (NCT02547987 and NCT02124902), analyzed by the National Cancer Institute’s Clinical Proteomic Tumor Analysis Consortium (CPTAC), have investigated less intensive neoadjuvant regimens for TNBC patients consisting of platinum + taxane and excluding anthracyclines^10^. However, in ∼50% of cases, pCR is not achieved, and these individuals experience dramatically shorter relapse-free and overall survival^11^. To further improve outcomes, clinical studies have also focused on intensifying therapy rather than de-escalating chemotherapy. Results from the KEYNOTE-522^12^ TNBC trial led to approval of adding an immune checkpoint inhibitor, pembrolizumab, to an already intensive chemotherapy backbone of carboplatin + paclitaxel followed by AC. However, addition of pembrolizumab improved the pCR rate by only ∼7% compared to the control arm^13^. Thus, identification of predictive markers for the efficacy of individual drugs would be useful in order to enhance response rates and to optimize and individualize treatments, as all chemotherapy agents have potentially life-threatening short- and long-term toxicities.

Our previous study identified weighted gene co-expression network analysis followed by Connect The Dots (WGCNA/CTD) informed mRNA-based biomarker panels that can predict differential response to single agent platinums and taxanes in multiple PDX and clinical datasets^14^. However, these predictive models were evaluated by leave-one-out cross-validation (LOOCV) without performance assessment of a fixed model on unseen, independent test data. Predictive biomarkers for the platinum + taxane combination were not identifiable. Furthermore, the mRNA-based biomarker panel did not provide insight into potential resistance mechanisms or therapeutic targets.

Here, we analyzed treatment response data to single agent carboplatin and docetaxel for a cohort of 50 TNBC PDXs derived from 46 different patients with TNBC (**Figure S1A**). Of these, we also generated combination response data for 42 PDXs to carboplatin + docetaxel combination. Treatment response data were integrated with baseline proteogenomic (DNA/mRNA/protein) profiles to perform a set of systematic analyses to address several key questions: 1) What are the molecular associates of response and resistance to single agent and combination treatments? 2) What genes and pathways are general features of chemotherapy resistance and responsiveness? and 3) What biomarker combinations are predictive of response to specific chemotherapy regimens?

## Results

### Combination carboplatin and docetaxel is largely ineffective at generating enhanced responses over the best single agent in TNBC PDXs

Patients with TNBC frequently receive combination chemotherapy treatments based on clinical data suggesting improved response from combination chemotherapy compared with single-agent chemotherapy^2,3^. However, averaging effects observed in group level data from clinical trials may not be applicable to individual patients^15^. Further, in clinical investigations, it is impossible to assign one patient to more than one treatment arm to deconvolute response to individual agents versus combinations. We hypothesized that this limitation can potentially be addressed through preclinical trials using PDX models.

To test this hypothesis, and to exploit this unique opportunity, we leveraged our collection of data from 49 TNBC PDX models and one estrogen-independent, low estrogen receptor-expressing model (FCP699) (reported to us originally as a TNBC, and propagated without estrogen as such until ER immunostaining showed low levels of ER in some nuclei) aggregated from three separate preclinical trials (**Figure S1B** and **Table S1A-B**). These in vivo preclinical trials included treatment arms consisting of control (no treatment) versus four weekly cycles of human-equivalent doses of carboplatin (50 mg/kg), docetaxel (20 mg/kg) or the combination at these doses^14^. Treatment effects were evaluated by log_2_ fold-change in tumor volume after four weeks vs. baseline and compared using general linear models (**Methods** and **Table S1B-D**). Quantitative and qualitative best clinical responses (**Methods**) to each treatment were summarized for each PDX model (**Figure 1A**). All PDXs had treatment responses documented for single agent carboplatin and docetaxel, while 84% (42/50) of PDXs had combination response information.

**Figure 1:**
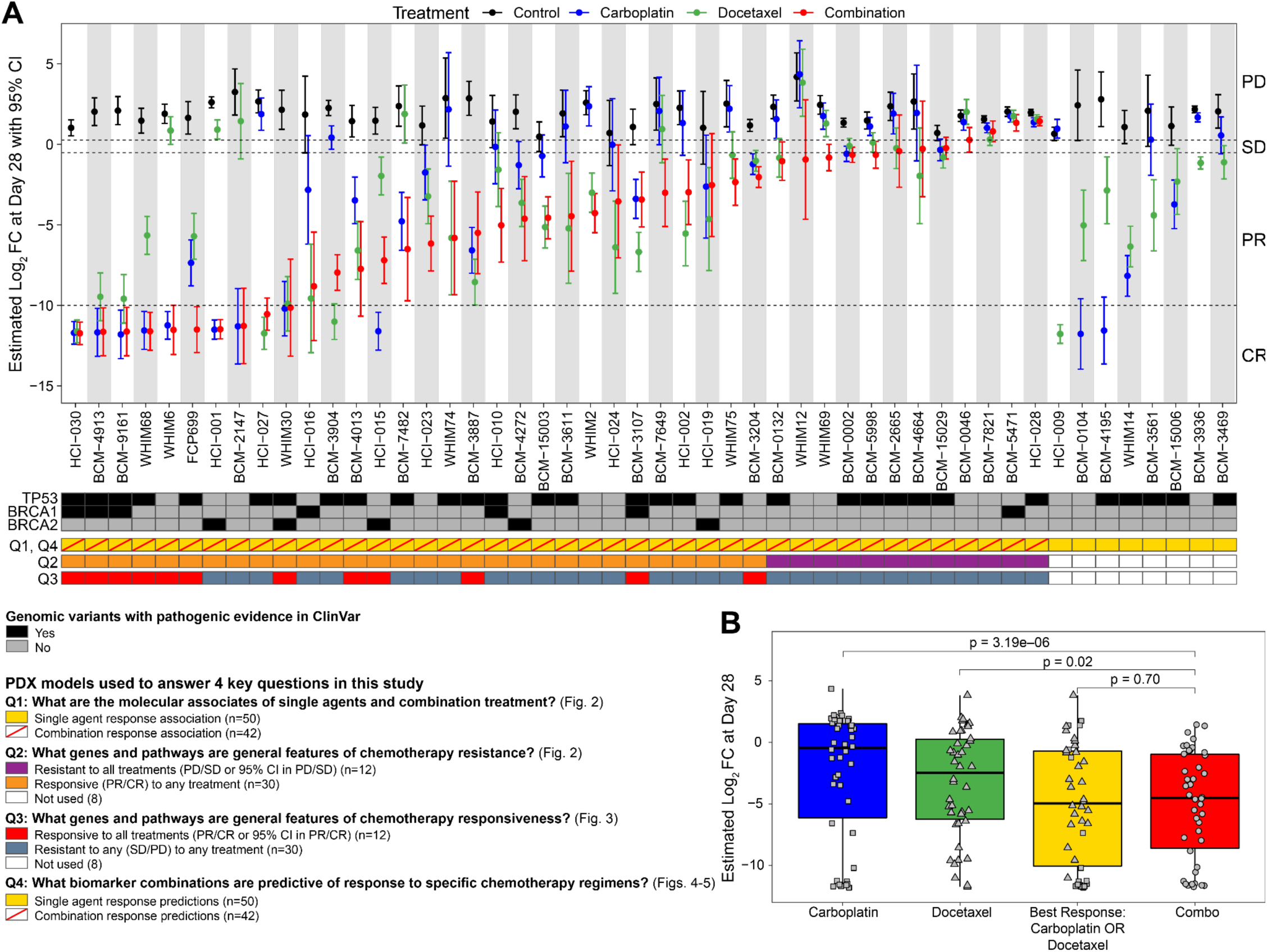
Combination carboplatin and docetaxel is largely ineffective at generating enhanced responses over the best single agent in TNBC PDXs. **A)** (Top panel) Quantitative tumor responses to chemotherapy in 50 TNBC PDX models comprising the “chemotherapy response cohort”. A general linear model for each PDX was generated to estimate mean log_2_ fold change (FC) in tumor volume at Day 28 vs Day 0 (baseline) and its associated 95% confidence interval (CI) for each treatment within a PDX (n ≥ 3 per treatment arm). Response was also qualitatively assessed based on modified RECIST 1.1 classification (PD, Progressive Disease; SD, Stable Disease; PR, Partial Response; CR, Complete Response). (Middle panel) Annotation of PDXs harboring pathogenic variants of TP53, BRCA1, BRCA2. No pathogenic PALB2 variants were identified. (Bottom panel) Annotations depicting sample stratification for analyses to address four key questions (Q1 to Q4) asked in this study. **B)** For PDXs with response to all treatment arms, boxplots depict quantitative chemotherapy responses for each PDX to single agent carboplatin (blue) and docetaxel (green), the best response to either single agent carboplatin OR docetaxel (yellow), and to the combination (red). These results suggest that combination treatment does not enhance tumor shrinkage over the best single-agent in TNBC PDXs. Boxes depict the interquartile range (IQR) of the scores with horizontal lines depicting the median. Whiskers extend to 1.5 x IQR from Q1 (25th percentile) and Q3 (75th percentile), respectively. P-values derived from paired Wilcoxon signed- rank tests.

Among PDXs that had treatment response information for all three treatment arms (**Figure 1A**, Q1), 71% (30/42) were responsive (complete response [CR] or partially responsive [PR]) to at least one treatment arm, while 29% (12/42) were resistant (stable disease [SD] or progressive disease [PD]) to all treatment arms, according to our modified RECIST criteria (mRECIST)^16,17^. When including the full cohort of 50 PDXs, 76% (38/50) were responsive to any treatment. When comparing quantitative tumor responses based on tumor volume change, single agent docetaxel led to significantly better tumor shrinkage in a greater number of PDX models compared to carboplatin at the doses evaluated (*n* = 18 vs *n* = 11, respectively, **Figure 1A** and **Table S1C**).

As expected, overall average tumor shrinkage was significantly greater in the combination arm compared to either the carboplatin (Wilcoxon signed-rank p = 3.19e−06) or docetaxel arms (Wilcoxon signed-rank p = 0.02) (**Figure 1B**). However, average shrinkage in the combination arm was generally comparable to the best single agent response for each PDX (p = 0.70) (**Figure 1B**). At the individual PDX level, combination treatment produced significantly enhanced responses over the best single agent in only 13% (4/30) of PDXs, was significantly worse than the best single agent in 12% (5/42), and did not qualitatively improve response assessed by mRECIST in 97% (29/30) of PDXs that had a response to any of the three arms tested (**Figure 1A**). This argues against a uniform additive benefit and highlights substantial response heterogeneity across PDXs. To evaluate whether the combination achieved greater than expected efficacy through independent contributions from both drugs, we assessed tumor responses using a Bliss independence framework^18^ (**Methods**). Under this model, the expected combination effect reflects the activity predicted if carboplatin and docetaxel act independently within the same tumor. At the cohort level, observed combination responses were significantly less inhibitory than predicted under independent drug action (paired t-test p = 1.7e−04, Wilcoxon signed-rank p = 8.3e−05, **Figure S1C**). When each PDX was evaluated individually, only one exhibited a combination response significantly greater than the Bliss prediction (**Figure S1D**). Together, these results indicate that additional benefit from carboplatin + docetaxel beyond what is achievable with a single agent is rare in this TNBC PDX cohort and further suggest that combination (versus single agent) chemotherapy may be unnecessary and support an approach to match patients to the most effective single agent.

### Molecular associates of chemotherapy response

To identify molecular correlates of treatment response and resistance that might provide mechanistic insight, we examined associations between mutation, copy number, mRNA, and protein profiles in baseline PDX tumors (**Table S2**) with treatment responses (**Table S3A-S3O**). Molecular profiling by whole exome sequencing, RNA-Seq, and mass spectrometry-based proteomic profiling in these PDXs has been reported previously^14^ and is described in **Methods**. Significant associations were observed for 2 non-silent mutations, 429 mRNAs, and 81 proteins for carboplatin response; 5 non-silent mutations, 222 mRNAs, and 44 proteins for docetaxel response; and 8 non-silent mutations, 190 mRNAs, and 55 proteins for combination treatment response (p < 0.01, Methods, **Figure 2A**, **Figure S2,** and **Table S3P**). No significant associations were found with copy number data at p < 0.01. At a more relaxed p < 0.05 threshold, high-level amplification events in 39 genes associated with carboplatin response, 22 genes associated with docetaxel resistance, and none were associated with combination response (**Figure S2**). Many genes found to be associated with carboplatin resistance were previously implicated in platinum resistance in cancer^19^ (**Table S3Q**). As an example, deleterious BRCA2 variants (n = 7) were associated with increased carboplatin response (**Figure S2A**) but not to docetaxel, nor to the combination. All 7 BRCA2-mutant models were carboplatin responders (7/7 vs 18/43, 42%, of BRCA2-wild-type; Fisher’s exact p = 0.010). Deleterious BRCA1 variants (n = 10) showed the same trend but did not reach significance in this analysis (7/10, 70%, responders vs 45% in BRCA1-wild-type; Fisher’s exact p = 0.29; **Figure S2A**). Because both genes converge on homologous-recombination deficiency, we pooled all deleterious BRCA1/2 variants (n = 16), which yielded a significant association with carboplatin response (13/16, 81%, vs 12/34, 35%; Fisher’s exact p = 0.005), supporting an HRD-driven effect. Consistent with a genetic rather than signature-level basis for this association, the HRD-associated mutational signature SBS3 was not a significantly assciated with carboplatin response (**Figure S2A**), in keeping with the limited resolution of exome-based signature estimation.

**Figure 2.**
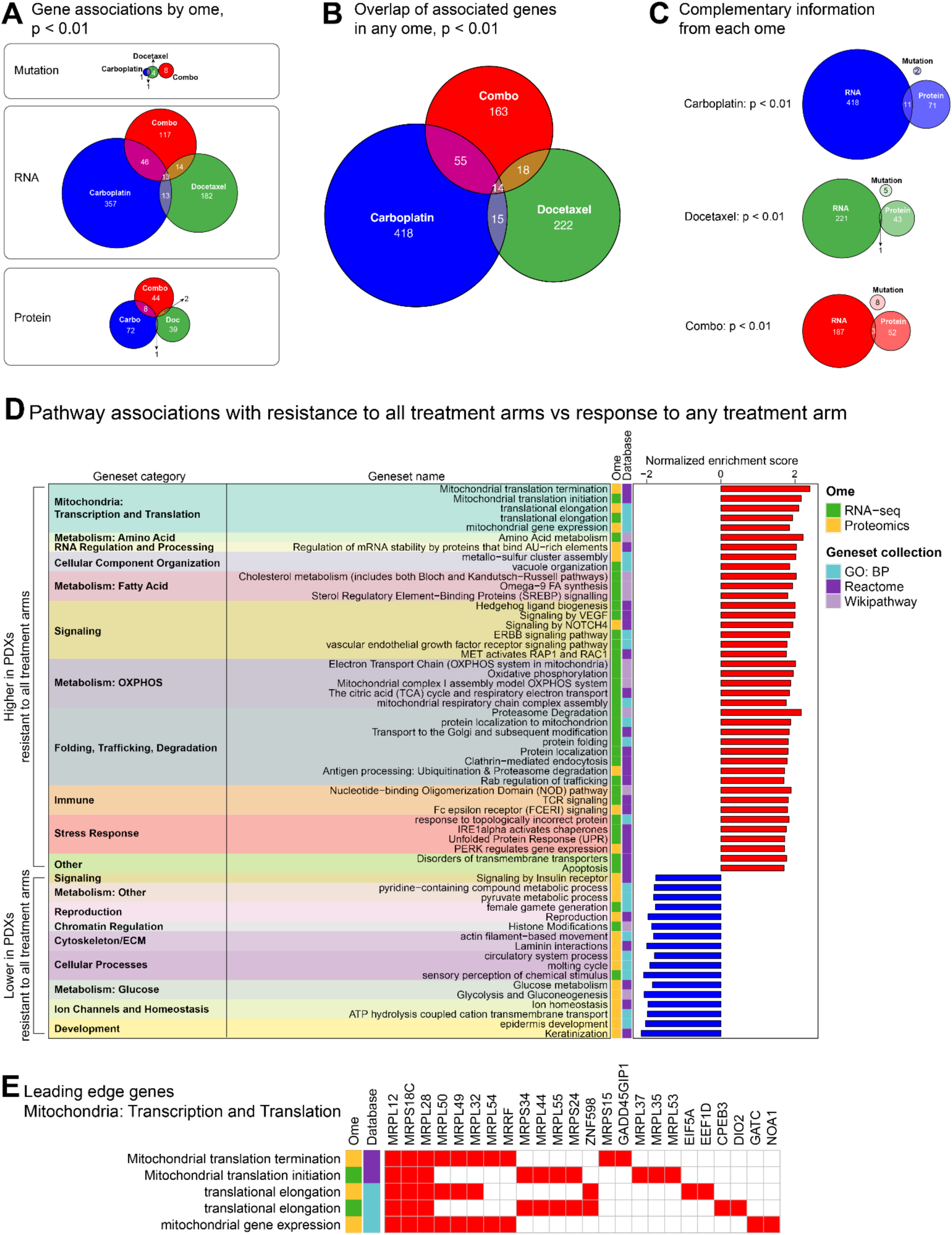
Molecular associates of chemotherapy response. **A)** Venn diagrams depicting overlap of treatment-associated genes from **Figure S2** (p < 0.01) by data type. **B)** Venn diagram depicting overlap of treatment-associated genes (p < 0.01) by any data type from **Figure S2**. **C)** Venn diagrams depicting overlap of significant (p < 0.01) molecular associations by treatment type. **D)** GSEA analysis with Gene Ontology: Biological Process (GO: BP), Reactome, and Wikipathway geneset collections using signed (indicating positive or negative sign of correlation coefficient) −log10 p-values from RNA-Seq and proteomics data in **Figure S3A**. Normalized enrichment scores of significant genesets (FDR < 0.05) are shown after applying weighted set cover to reduce geneset redundancy. Genesets were further categorized into categories by manual curation guided by leading edge genes in each geneset. **E)** Top 10 leading edge genes for mitochondrial transcription and translation pathways.

There were some associations that were shared between different treatments (**Figure 2B, Table S3R)**. For example, Glutamate Rich 6 (ERICH6) variants were associated with increased response to both single-agent carboplatin and docetaxel, but not the combination (**Figure S2A-C**). For the same treatment, different types of molecular data identified distinct sets of associated genes (**Figure 2C, Table S3S**), each contributing unique yet complementary information. This finding is not unexpected given that different types of molecular data are not fully correlated with one another. For example, in this study, the median gene-wise Spearman’s correlation between mRNA and protein was 0.46, consistent with the poor mRNA-protein correlation observed for many genes in other proteogenomic studies^20^. This underscores the importance of multi-omic analyses to gain a comprehensive understanding of the genes that may underlie treatment response.

### Molecular features associated with lack of chemotherapy response

In PDXs that were treated with all 3 treatment arms, at least one treatment induced tumor shrinkage for most models, yet 29% (12/42) of PDXs were resistant to all chemotherapy regimens tested (**Figure 1A**, Q2). To identify molecular associations with lack of any chemotherapy response vs response to any treatment, we compared PDXs without a response to all treatment arms vs PDXs with tumor shrinkage due to any regimen using data generated across all omic platforms (**Figure S3A** and **Table S4A-D**). In PDXs resistant to all treatments, RNA-Seq and protein data associated gene sets broadly related cellular respiration and metabolism, including mitochondrial transcription and translation, OXPHOS, and fatty acid and amino acid metabolism to resistance (**Figures 2D-E, Figures S3B-C,** and **Table S4E-J**). Upregulation of OXPHOS and fatty acid metabolism pathways were also found to be associated with tumors resistant to neoadjuvant carboplatin + docetaxel in the CPTAC-TNBC clinical trial^10^. Moreover, targeting fatty acid pathways and subsequent mitochondrial function has been shown to inhibit TNBC^21^. Of note, multiple proteostatic pathways related to protein folding, trafficking, and degradation along with unfolded protein response (UPR) pathways related to endoplasmic reticulum stress were heightened in PDXs resistant to all chemotherapy arms (**Figure 2D** and **Figure S3D**) while chromatin regulation and glucose metabolism were downregulated in these PDXs (**Figure 2D** and **Figure S3E**).

### KRT5 is a chemotherapy response marker for carboplatin, docetaxel, and their combination

In contrast, in PDXs responsive to any treatment arm, basal marker proteins and structural components of cells, such as cytokeratins KRT5, KRT6B, and KRT17 were among the most significantly upregulated genes especially at the protein level (**Figure S3A**). These gene level results support pathway associations when comparing PDXs with response to all treatment arms compared to those with lack of response to any regimen (**Figure 1A**, Q3) where cytoskeletal/ECM gene sets were found to be higher in PDXs with response to all treatment arms (**Figure 3A**) (and also lower in PDXs resistant to all treatment arms (**Figure 2D**). To test whether these PDX-derived markers translate to human disease, we examined baseline tumors from the CPTAC-TNBC neoadjuvant chemotherapy trial^10^, the closest available human counterpart to our PDX treatment and molecular profiling design. Since cytokeratin upregulation in the PDXs was observed by a subset of responsive models rather than a uniform shift (**Figure 3C**), we hypothesized that any corresponding signal in CPTAC-TNBC could also be confined to a subset of tumors. Since such subset-restricted differences are not well captured by tests of central tendency (e.g. Wilcoxon rank-sum test and t-test), we applied the Anderson-Darling (AD) test, which is more sensitive to differential expression in distribution tails. We previously applied this approach to detect tumor-associated antigens with subset-restricted expression in pan-cancer data^20^. Of the three markers, only KRT5 was significantly different at the RNA level by AD, with protein data nearing p < 0.05. Wilcoxon rank-sum comparisons were not significant for any of the three. The AD result is consistent with the PDX data and indicates that a subset of non-pCR tumors have lower KRT5 than pCR tumors (**Figure 3D**). Both KRT5 mRNA and protein abundance were able to discriminate non-pCR CPTAC-TNBC samples from pCR samples with an AUROC of 0.69 and 0.65, respectively (**Figure 3E**). Importantly, both AUROC curves demonstrated appreciable sensitivity with a 0% false positive rate, indicating high specificity of KRT5 to distinguish non-pCR tumors to neoadjuvant chemotherapy. We further validated KRT5 by IHC staining of baseline tumors prior to treatment from our PDX cohort (**Figure 3F**) where staining intensity and frequency were quantified to generate an Allred score (**Figure 3G**). By using an Allred score, KRT5 quantification achieved an AUROC of 0.83 in discriminating PDX tumors responsive to any chemotherapy arm in our cohort (**Figure 3H**), providing orthogonal validation and supporting clinical translatability of using KRT5 as an IHC-based biomarker of responsiveness to non-anthracycline-based chemotherapy regimens consisting of carboplatin, docetaxel, or the combination.

**Figure 3.**
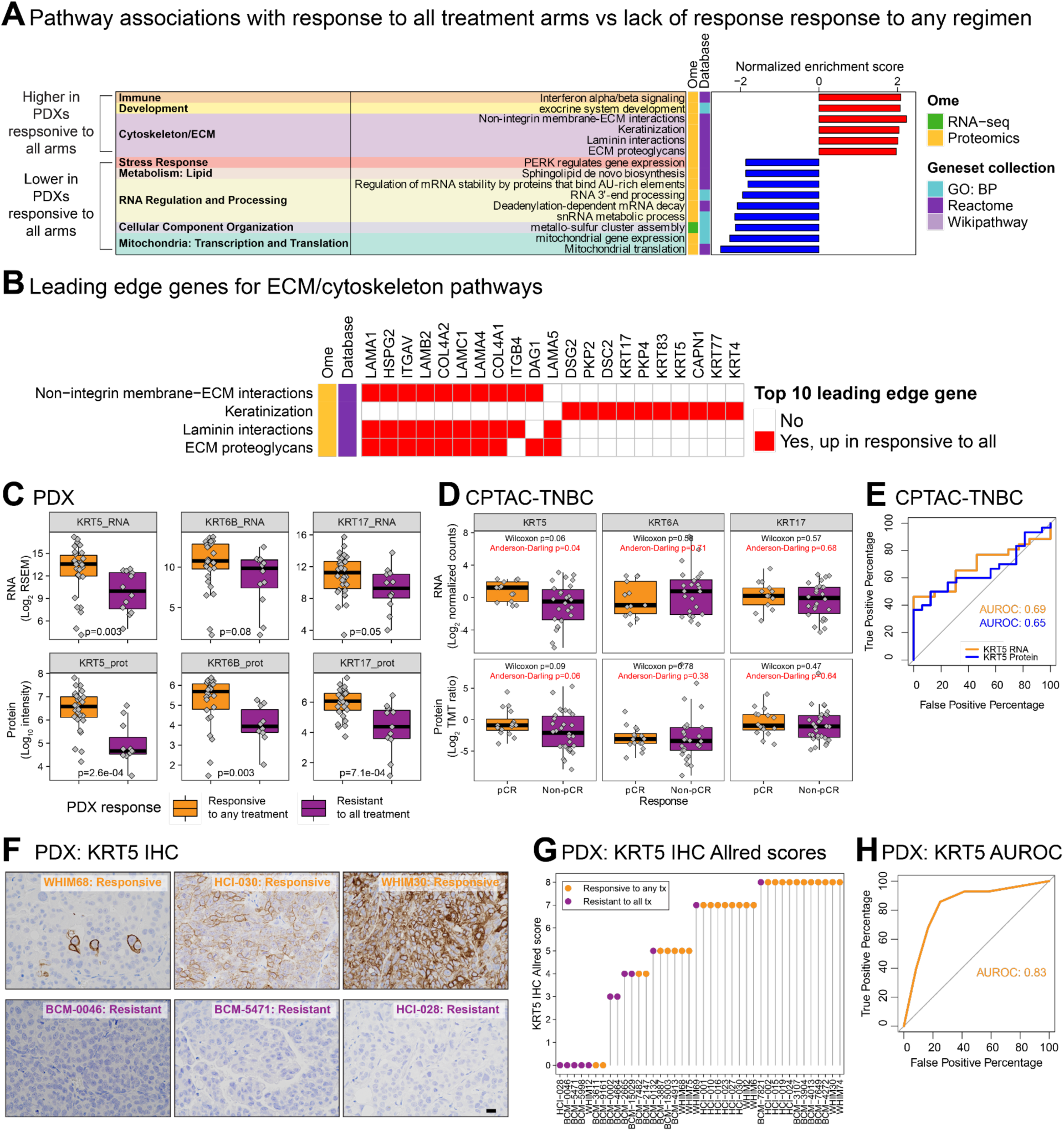
KRT5 is a general marker of chemotherapy responsiveness. **A)** GSEA analysis with Gene Ontology: Biological Process (GO: BP), Reactome, and Wikipathway geneset collections using signed (indicating positive or negative sign of correlation coefficient) −log10 p-values from RNA-Seq and proteomics data. **B)** Top 10 leading edge genes from cytoskeleton/ECM pathways. **C)** Boxplots showing basal cytokeratin genes that are higher in PDXs responsive to any treatment vs PDXs resistant to all treatments. **D)** Similar to (C) except showing the same cytokeratin genes in the CPTAC-TNBC dataset comparing pCR and non-pCR patients. For (C-D), boxes depict the interquartile range (IQR) of the scores with horizontal lines depicting the median. Whiskers extend to 1.5 x IQR from Q1 (25th percentile) and Q3 (75th percentile), respectively. P-values derived from Wilcoxon rank-sum tests. **E)** Plot depicting AUROC using KRT5 RNA or protein levels from CPTAC-TNBC study^10^ to discriminate non-pCR tumors from pCR tumors. **F)** Representative images of KRT5 IHC in baseline PDX tumors prior to treatment. Scale bar = 10 µm. **G)** Plot showing Allred IHC scores for KRT5 quantified from IHC images. **H)** AUROC using KRT5 Allred scores in predicting response to any chemotherapy treatment arm for TNBC PDXs in this study.

### Predicting objective response, complete response, and pathologic complete response

Following the identification of individual genes associated with differential chemotherapy responses (**Figure 2**) machine learning models were next employed to identify biomarker combinations predictive of complete response/pathologic complete response (pCR) and complete response plus partial response (CR/PR) or objective response to chemotherapy, both clinically meaningful response cutoffs. Individual datasets from this study, and previously described external resources (**Table 1**), were used to train separate logistic regression models using a state-of-the-art feature selection process using the Protein Marker Selection (ProMS) tool we previously developed^22^. ProMS identifies a group of minimally redundant, and therefore complementary, markers associated with an attribute label, which for this study is chemotherapy response, where each marker represents a biological function or pathway defined by co-expressed genes. Although initially developed for protein marker selection using single or multi-omics data, ProMS can also be implemented for RNA marker selection, which is how it was applied in this study. ProMS selected features to predict CR/pCR for platinum, taxane, and their combination, respectively, using transcriptomics profiling data (**Figure S4A** and **Methods**). Due to the small sample size of each dataset, five features were selected for model training. For each treatment type, the largest available clinical dataset was reserved as the independent test dataset for final model evaluation. Before model development, we evaluated the predictive utility of each remaining dataset using set-aside cross-validation within that dataset. Prediction performance of models trained using single datasets were highly variable and close to random (0.50) as measured by area under ROC curve (AUROC) in the set-aside cross-validation data from the same dataset (**Figure S4B**). In particular, the taxane Dataset 07 from Savage et al, 2020^23^ had a low AUROC of 0.35 and was omitted in subsequent analyses.

**Table 1.** TNBC Datasets Used For Chemotheropy Reponse Prediction.

| No | PMID/study name | Platform | Sample | Therapy received before response assessment | Response metric | Total samples | Responsive / Resistant | Role<sup>§</sup>: Figure 4 / Figure 5 |
| --- | --- | --- | --- | --- | --- | --- | --- | --- |
| Platinum datasets |  |  |  |  |  |  |  |  |
| 01 | This study | RNA-Seq | PDX | Carboplatin | Responsive: CR; Resistant: PR,SD,PD | 50 | 11 / 39 | Train / Train |
|  |  |  |  |  | Responsive: CR,PR; Resistant: SD,PD | 50 | 25 / 25 |  |
| 02 | PMID: 32546838<br>Savage et al., 2020 | RNA-Seq | PDX | Cisplatin | Responsive: CR; Resistant: PR,SD,PD | 20 | 7 / 13 | Train / Train |
|  |  |  |  |  | Responsive: CR,PR; Resistant: SD,PD | 20 | 12 / 8 |  |
| 03 | PMID: 37029129<br>Brugge et al., 2023 | RNA-Seq | PDX | Cisplatin | Responsive: CR; Resistant: PR,SD,PD | 39 | 8 / 31 | Train / Train |
|  |  |  |  |  | Responsive: CR,PR; Resistant: SD,PD | 39 | 12 / 27 |  |
| 04 | PMID: 20100965<br>DFCI | Microarray | Human tumors | Cisplatin | Responsive: CR; Resistant: PR,SD,PD | 24 | 3 / 21 | Test / Train |
|  |  |  |  |  | Responsive: CR,PR; Resistant: SD,PD | 24 | 15 / 9 |  |
| 05 | This study<sup>†</sup> | RNA-Seq | PDX | Carboplatin | Responsive: CR; Resistant: PR,SD,PD | 11 | 1 / 10 | Not used / Test |
| Taxane datasets |  |  |  |  |  |  |  |  |
| 06 | This study | RNA-Seq | PDX | Docetaxel | Responsive: CR; Resistant: PR,SD,PD | 50 | 4 / 46 | Train / Train |
|  |  |  |  |  | Responsive: CR,PR; Resistant: SD,PD | 50 | 36 / 14 |  |
| 07 | PMID: 32546838<br>Savage et al., 2020 | RNA-Seq | PDX | Paclitaxel | Responsive: CR; Resistant: PR,SD,PD | 20 | 8 / 12 | Omitted* |
|  |  |  |  |  | Responsive: CR,PR; Resistant: SD,PD | 20 | 13 / 7 |  |
| 08 | PMID: 24669015<br>MDA | RNA-Seq | Human tumors | Paclitaxel<sup>‡</sup> | Responsive: CR; Resistant: PR,SD,PD | 22 | 3 / 19 | Train / Train |
|  |  |  |  |  | Responsive: CR,PR; Resistant: SD,PD | 22 | 7 / 15 |  |
| 09 | PMID: 35623341<br>ISPY2 | Microarray | Human tumors | Paclitaxel<sup>‡</sup> | Responsive: CR; Resistant: PR,SD,PD | 132 | 19 / 113 | Test / Train |
|  |  |  |  |  | Responsive: CR,PR; Resistant: SD,PD | 132 | 80 / 52 |  |
| 10 | PMID: 28376176<br>BEAUTY | RNA-Seq | Human tumors | Paclitaxel<sup>‡</sup> | Responsive: CR; Resistant: PR,SD | 39 | 11 / 28 | Not used / Test |
| Platinum + taxane datasets |  |  |  |  |  |  |  |  |
| 11 | This study | RNA-Seq & proteomics<sup>#</sup> | PDX | Carboplatin + docetaxel | Responsive: CR; Resistant: PR,SD,PD | 42 | 10 / 32 | Train / Train |
| 12 | PMID: 36001024<br>CPTAC-TNBC | RNA-Seq & proteomics<sup>#</sup> | Human tumors | Carboplatin + docetaxel | Responsive: pCR; Resistant: Non-pCR | 39 | 13 / 26 | Train / Train |
| 13 | PMID: 35857626<br>COH-TNBC | RNA-Seq | Human tumors | Carboplatin + paclitaxel | Responsive: pCR; Resistant: Non-pCR | 41 | 21 / 20 | Test / Train |
| 14 | This study<sup>†</sup> | RNA-Seq | PDX | Carboplatin + docetaxel | Responsive: CR; Resistant: PR,SD,PD | 11 | 1 / 10 | Not used / Test |

\*Dataset 07 omitted due to low prediction performance as a single dataset (AUROC = 0.35)
<sup>†</sup>11 additional PDXs not part of the original 50-model chemotherapy response cohort
<sup>‡</sup>Response assessment derived from MRI-based measurements after administration of paclitaxel but prior to AC
<sup>#</sup>Proteomics data used together with RNA-Seq for protein-informed RNA marker selection via ProMS<sup>mo</sup> (**Figure S5**)
<sup>§</sup>Datasets used as independent tests in initial models (**Figure 4**) were incorporated into composite model training (**Figure 5**) to maximize sample size

Since models trained on individual datasets were unstable and not predictive, likely due to interpatient heterogeneity in chemotherapy resistance pathways, which cannot be fully captured by small sample sizes, datasets of the same treatment type were batch corrected and merged to increase sample size (**Figures S4C- F**). Within each merged dataset, the largest clinical cohort was designated as the independent test set (**Table 1**, Role: Figure 4 column). Beyond batch correction, which does not include response information, these held-out datasets were not used for feature selection or model fitting. They were reserved exclusively for evaluating fully trained models. Since our previous study showed that prediction performance was highly dependent on the feature selection method^14^, we compared four different feature selection methods: (1) genes associated with quantitative responses (regression genes), (2) WGCNA-CTD, (3) ProMS, and (4) CR/PR-associated genes, to evaluate their ability to predict objective (CR/PR), complete, and pathologic complete responses (**Methods**). Specifically, ProMS optimization identified 10, and 9 features for platinum, and taxane CR/PR predictors, respectively, and 5, 6, and 10 features for the platinum, taxane, and combination (p)CR predictors (**Figure S5**). Selected features for predicting objective responses for platinum and taxane from each of the feature selection methods were largely non-overlapping (**Figure 4A**, and **Table S5**). However, many of these genes were functionally related. Multiple methods selected different zinc-finger transcriptional regulators such as ZFP37, ZNF256, ZNF394, and ZNF462 for platinum, and ZNF85, ZNF100, ZNF493, ZNF322, and ZNF669 for taxane. This suggests that each of the feature selection methods may capture both shared and distinct aspects of chemotherapy response biology. Logistic regression models trained using features from each method showed variable performance in predicting objective CR/PR responses to platinum and taxane chemotherapy in the set-aside, independent test data (**Figure 4B**), ranging from 0.55 - 0.74 and 0.52 - 0.58 AUROC, respectively. For single-agent platinum regimens, pooling all non-overlapping features from each method resulted in higher predictive performance (AUROC = 0.76) than any individual feature selection method alone, though this pooling strategy did not improve performance for taxane regimens (**Figure 4B**).

**Figure 4.**
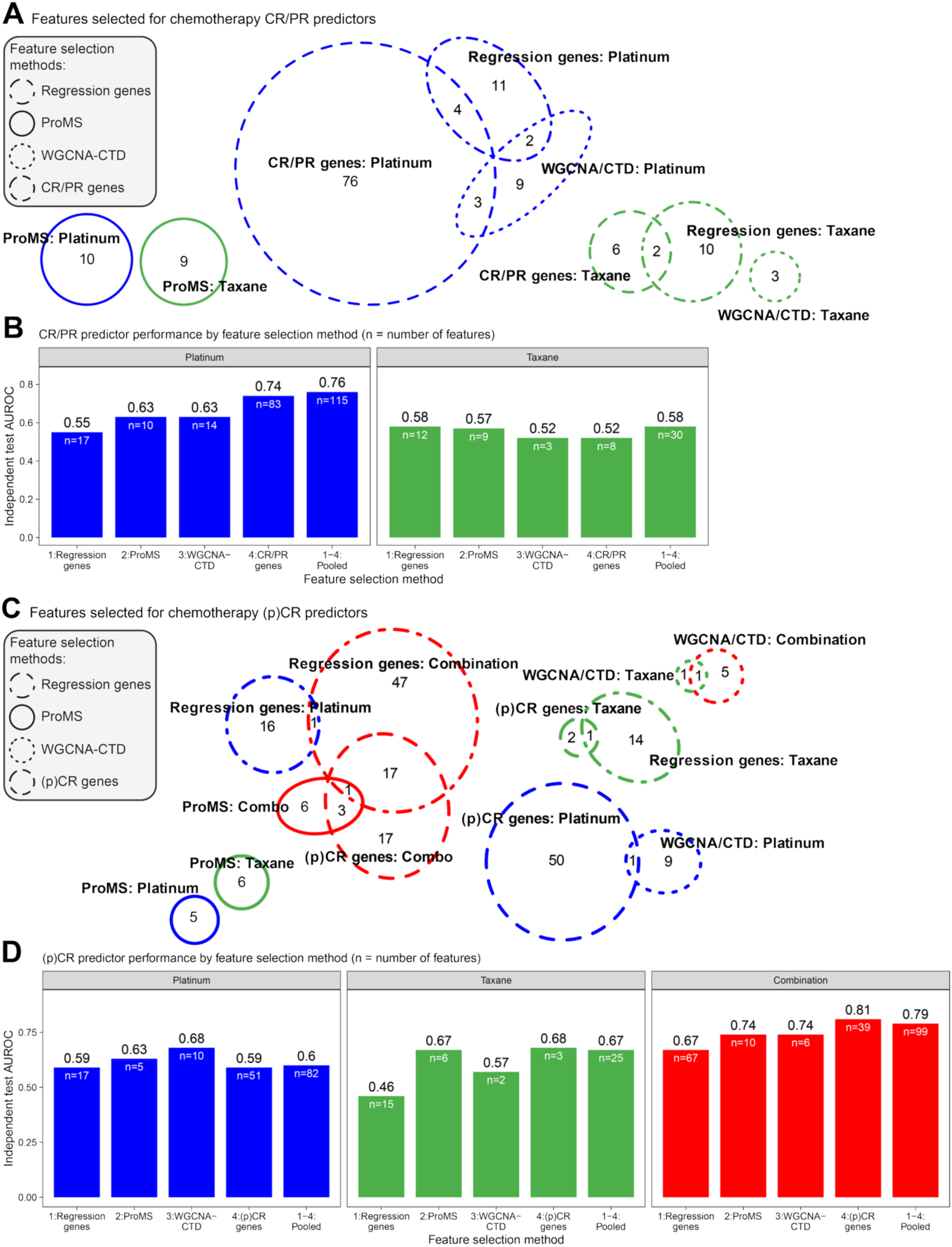
Predicting objective response, complete response and pathologic complete response. **A)** Venn diagram showing overlapping features of regression genes, WGCNA/CTD, ProMS, and CR/PR-associated genes for predicting objective response (CR/PR response). **B)** AUROC performance on independent test data of logistic regression models trained to predict CR/PR using features selected by different feature selection methods as indicated or using pooled features. n = number of features. **C)** Same as (A) but used for predicting complete/pathologic complete response. **D)** Same as in (B) except showing AUROC performance on independent test data of models trained to predict complete/pathologic complete response. n = number of features.

For complete and pathologic complete response predictors, the same four feature selection methods again identified largely distinct sets of genes (**Figure 4C** and **Table S6**) similar to the patterns observed for the CR/PR models. Logistic regression models trained using these feature sets also showed variable performance across chemotherapy regimens (**Figure 4D**). For platinum, WGCNA-CTD features generated the highest performance predictor with an AUROC of 0.68 (**Figure 4D**). For taxane and combination treatments, (p)CR associated genes generated the highest performance predictor with an AUROC of 0.68 and 0.81, respectively (**Figure 4D**). These results indicate that, as with the CR/PR predictor, the choice of feature selection methods influences (p)CR prediction performance. Since the number of candidate features differed across methods, these comparisons reflect both the selection strategy and feature-set size.

We next examined whether incorporating protein data could further enhance prediction. The ProMS method is unique for its ability to be extended to the multi-omics setting, leveraging multi-omic data for biomarker discovery within a single omic layer of interest^22^. For the platinum + taxane predictor, both RNA and protein data were available for training datasets. Since 10 biomarkers achieved the best cross-validated performance by using RNA data alone (**Figure S5G**), merged RNA and protein data (**Figure S4E-F**) were used together to also select a group of 10 protein-informed, RNA biomarkers by running ProMS in multi-omics mode (ProMS^mo^). Interestingly, only one of these genes, LGALS1, overlapped with the RNA-only informed ProMS model (**Figure S5H** and **Table S6M-N**). Performance of the platinum + taxane predictor trained with features selected by ProMS^mo^ (RNA + protein) outperformed ProMS (RNA) on independent test data, achieving an AUROC of 0.85 (**Figure S5I**). These results suggest protein data heavily impact RNA marker selection and can enhance prediction performance when used together with RNA data. It is notable that the protein-informed model achieved a 0% false positive rate with close to 60% sensitivity, as revealed by the sharp initial climb in the AUROC curve, underscoring its high specificity in predicting pCR tumors in response to platinum + taxane chemotherapy.

In sum, these results highlight the potential of using machine learning models, and the integration of protein data when available, to select TNBC tumors that may respond to either single agent carboplatin, docetaxel or their combination, consistent with our previous work^14^.

### Optimizing feature selection approaches improves prediction of chemotherapy response

To improve upon the transcriptome-based models and to generate predictive models that can be taken forward in future PDX-based studies and clinical trials, final, composite, logistic regression models for prediction of platinum, taxane, and platinum + taxane clinical/pathologic complete response were generated by incorporating the clinical datasets previously held out as independent tests in **Figure 4** into the training set (see **Table 1**, Role: Figure 5 column). These composite models represent “the best that can be done” until additional datasets become available. The number of features by each method was determined using the same criteria applied in **Figure 4** (**Methods**). The four feature selection approaches again identified distinct sets of genes (**Figure 5A** and **Table S7**). Performance of logistic regression models to predict CR/pCR was assessed using Monte-Carlo cross-validation (MCCV) where 70% of the data was randomly used for training and the remaining 30% was held out for testing. This process was repeated 50 times to approximate how they would perform on unseen samples (**Figure 5B**). This approach allowed us to use the full dataset for training while still obtaining an internal estimate of generalizability. Results of this process generated higher performing models than previously described with the composite platinum predictor reaching a mean AUROC of 0.83 using CR-associated genes and pooled features, the composite taxane predictor achieving a mean AUROC of 0.81 with regression and CR-associated genes, and the composite platinum + taxane predictor achieving a mean AUROC of 0.86 with CR-associated genes (**Figure 5B**). By using pooled features from the composite models, logistic trained predictors showed extremely high performance when applied to the models used to train them as expected (**Figure 5C**). These logistic trained models using pooled composite features were then applied to independent datasets not used at any training stage. For the taxane predictor, the BEAUTY clinical trial^24^, in which MRI-based responses were assessed after TNBC patients received paclitaxel only, served as the independent test set (AUROC = 0.86). For the platinum and platinum + taxane predictors, an additional cohort of 11 PDXs not part of the original 50 from **Figure 1** served as the independent test set (AUROC = 0.80 for both) (**Figure 5D**). Further performance evaluation of these composite predictors await further evaluation in datasets that may become available in the near future, including data from TBCRC 030^25^ and RESPONSE (NCT05020860) clinical trials, or in future experimental settings using PDX/PDX-derived organoids.

**Figure 5.**
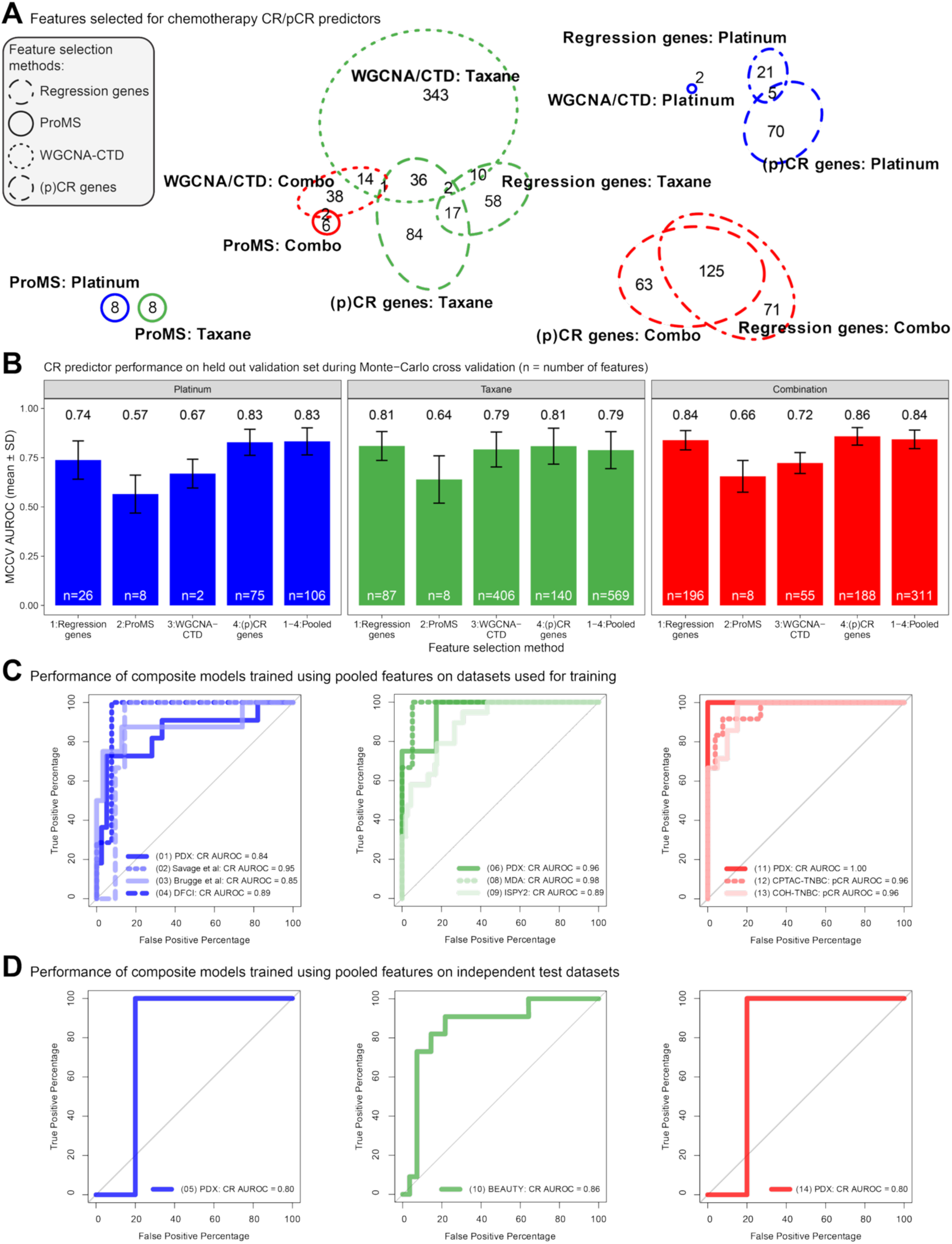
Optimizing chemotherapy complete/pathologic complete response predictors. **A)** Venn diagram showing overlap of features selected by regression-associated genes, ProMS, WGCNA/CTD, and for (p)CR- associated genes for predictors using all possible datasets. **B)** Mean of Monte-Carlo cross validation AUROC performance on held-out dataset of composite logistic regression models trained to predict complete/pathologic complete response (repeated 50 times) using features selected by different feature selection methods as indicated or using pooled features. n = number of features. **C)** AUROC curves showing performance of composite models trained on pooled features when applied to datasets used for training. Number in parenthesis indicate dataset from **Table 1**. **D)** AUROC curves showing performance on individual datasets not used for training that includes PDX models and external patient cohorts. Number in parenthesis indicate dataset from **Table 1**.

Beyond their predictive performance, we asked what biology these predictors capture by characterizing them systematically across all the models rather than highlighting individual genes. Over-representation analysis of the pooled features for each treatment context (**Figure S6A**) was enriched with mechanistically coherent pathways by treatment. Platinum predictors mapped to the Fanconi anemia DNA-crosslink-repair pathway (BRIP1, FANCI, POLH, TOP3A, USP1; **Figure S6B**), consistent with the HRD-driven carboplatin association (**Figure 2**). Taxane predictors were dominated by extracellular-matrix and cell-adhesion programs such as collagens and laminins, along with FN1, THBS1 and ITGB5, together with growth-factor molecules (EGFR, FGFR1, PDGFA/B) acting through Rap1 and PI3K-Akt signaling (**Figure S6B**), in line with the cytoskeletal/ECM programs of responsive tumors (**Figure 3**). Combination predictors were enriched for mitotic cell-cycle and interferon/cytokine immune signaling (STAT1, IRF9, OAS1-3, IFIT1, SOCS1/3; **Figure S6B**). Annotating the gene-level composition of these pathways by selection method (**Figure S6B**) showed that pathway enrichment arose through two complementary patterns. First, different methods contributed distinct genes to the same pathway, as exemplified by the Fanconi anemia pathway, where BRIP1, FANCI, and USP1 were nominated by regression, ProMS, and (p)CR-associated gene analyses, respectively. Second, the same gene was selected independently by multiple methods, most notably OAS2, which was identified by all four approaches, while STAT1, OAS3, and SOCS1/3 were identified by two or three methods.

To illustrate the clinical utility of our predictors, we derived candidate score cutpoints from the independent test data using two complementary strategies. A screening cutpoint favoring sensitivity, to rule out non-responders, and a diagnostic cutpoint at the sensitivity-specificity crossover (**Table 2** and **Methods**). We did not perform a comparable analysis for the platinum predictor because the validation cohort contained insufficient numbers of responders and non-responders to support reliable estimation of clinically meaningful cutpoints. For the taxane predictor evaluated in BEAUTY, the screening cutpoint identified responders with 90.9% sensitivity and 78.6% specificity, while the diagnostic cutpoint was balanced at 81.8% sensitivity and 85.7% specificity (**Table 2**). For the protein-informed platinum + taxane predictor (ProMS^mo^), the screening cutpoint captured 95.2% of responders but at lower specificity (40.0%), whereas the diagnostic cutpoint was balanced at 76.2% sensitivity and 75.0% specificity (**Table 2**). Consistent with its steep AUROC curve, more stringent thresholds can reach 100% specificity (**Figure S5I**), identifying a subset of pCR tumors with complete (100%) positive predictive value. These cutpoints are intended as illustrative decision rules requiring prospective validation, but they demonstrate that a single predictive score can be tuned to rule tumors either in or out of platinum- and taxane-based chemotherapy.

**Table 2.** Candidate classification cutpoints for the chemotherapy-response predictors.

| Cutpoint | Performance score | Response |  | Sensitivity (95% CI) | Specificity (95% CI) | PPV |
| --- | --- | --- | --- | --- | --- | --- |
|  |  | (p)CR | Non-(p)CR |  |  |  |
| Taxane predictor, composite features: |  |  |  |  |  |  |
| BEAUTY independent test; AUROC 0.86; n = 39 (11 CR, 28 non-CR) |  |  |  |  |  |  |
| Screening | > 0.007 | 10 | 6 | 90.9% (58.7-99.8) | 78.6% (59.1-91.7) | 62.5% |
|  | ≤ 0.007 | 1 | 22 |  |  |  |
| Diagnostic | > 0.0107 | 9 | 4 | 81.8% (48.2-97.7) | 85.7% (76.3-96.0) | 69.2% |
|  | ≤ 0.0107 | 2 | 24 |  |  |  |
| Platinum + taxane predictor, ProMS<sup>mo</sup> features: |  |  |  |  |  |  |
| COH-TNBC independent test; AUROC 0.85; n = 41 (21 pCR, 20 non-pCR) |  |  |  |  |  |  |
| Screening | > 0.131 | 20 | 12 | 95.2% (76.2-99.9) | 40.0% (19.1-63.9) | 62.5% |
|  | ≤ 0.131 | 1 | 8 |  |  |  |
| Diagnostic | > 0.232 | 16 | 5 | 76.2% (52.8-91.8) | 75.0% (50.9-91.3) | 76.2% |
|  | ≤ 0.232 | 5 | 15 |  |  |  |

For each predictor a screening cutpoint (favoring sensitivity, for ruling out non-responders) and a diagnostic cutpoint (at the sensitivity-specificity crossover) are shown. Counts are tumors above / below the score cutpoint cross-classified by observed response (CR, complete/pathologic complete response). The screening cutpoint is the highest sensitivity off the tail of the ROC curve with the maximal specificity among those candidates; the diagnostic cutpoint is the score at which the sensitivity and specificity curves cross.
CI, exact (Clopper-Pearson) 95% confidence interval; PPV, positive predictive value (percentage of predicted-sensitive tumors that responded). Response definitions differ between panels (BEAUTY: MRI-based CR on paclitaxel; ProMS<sup>mo</sup>: pCR on platinum + taxane).

In summary, these findings highlight the impact of different feature-selection strategies and the benefit of maximizing sample size to train high-performance models for predicting chemotherapy response, and show that the selected features, though largely distinct, converge on coherent, treatment-appropriate biological programs.

## Discussion

Cytotoxic chemotherapy regimens for TNBC patients frequently include several agents given in combination, or in series, without prior knowledge of whether a given agent would be effective or ineffective for an individual patient. Deconvolution of response to multiple single agents is difficult-to-impossible to address effectively in clinical trials as it is not possible to assign the same patient to multiple treatment arms. We addressed this critical issue by leveraging a clinically relevant, large cohort of 50 orthotopically transplanted PDXs where each model was assessed for response to human equivalent doses of single agent carboplatin, docetaxel, or the combination. We also leveraged multiple publicly available PDX and neoadjuvant TNBC datasets in which mice or patients were treated with similar chemotherapy regimens for validation throughout the study to suggest clinical relevance of our findings, particularly if these gene panels can be refined. Our investigation into the lack of enhanced combination response compared to the best single agent for a given TNBC PDXs has numerous important biological and clinical implications discussed below.

There is a great need to identify additional biomarkers for precision diagnostics beyond ER, PR, and HER2. In contrast to neoadjuvant pCR predictors for taxane-anthracycline based regimens using gene expression^26,27^, predictors for anthracycline-free and platinum-containing regimens are understudied. Incorporation of carboplatin with taxane-anthracycline regimens improved pCR rates compared with taxane-anthracycline alone in TNBC patients from the BrighTNess trial^6^ (58% vs 31%, respectively). Furthermore, multiple TNBC trials examining anthracycline-free regimens consisting only of carboplatin + taxane have shown pCR rates above 50%^10,28–30^, suggesting opportunities for chemotherapy optimization^31,32^. The ongoing SCARLET trial (NCT05929768), a phase III randomized study for TNBC patients, is now prospectively testing whether anthracycline-free taxane-platinum chemo-immunotherapy with pembrolizumab is non-inferior to anthracycline-containing regimens, further highlighting a focus toward optimization strategies for TNBC. However, even with the addition of pembrolizumab to carboplatin/taxane/anthracycline-containing chemotherapy, this regimen provided only a 7% improvement in pCR rate compared with chemotherapy alone^13^, suggesting a critical need for predictive biomarkers to identify patients that will benefit from immune-checkpoint inhibition and to determine whether anthracyclines can be safely omitted.

Since tumor shrinkage is a meaningful endpoint of neoadjuvant chemotherapy that leads to downstaging and improving surgical outcomes, our CR/PR predictors, especially for single agent platinum (**Figure 4A-B**) could also have implications for personalizing treatment strategies, optimizing patient selection for neoadjuvant therapy, and guiding clinical decision making to minimize overtreatment. The taxane CR/PR predictors did not perform as well as their CR counterparts which might be attributed to the inclusion of multiple CR/PR datasets with response annotation from MRI-based clinical measurements. Discrepancies might arise in generating accurate response labels, such as distinguishing PR from SD, and therefore may impact the consistency and reliability of the training data, leading to reduced model performance and less predictive power in borderline response categories.

The high performance and high specificity of our protein-informed, multi-omic pCR predictor for platinum + taxane (**Figure S5I**) has important implications for identifying patients that will respond to chemotherapy without the need for additional pembrolizumab and anthracycline/cyclophosphamide. Integrating proteomic data with transcriptomic data demonstrates the added value of incorporating diverse molecular profiles to enhance chemotherapy response predictions. Moreover, the ProMS and ProMS^mo^ selected biomarkers that comprise our chemotherapy response predictors to single-agent platinum, taxane, and their combination form a relatively small set of genes that is tractable for further validation and clinical translation and therefore attractive for future development into a precision diagnostic. We have trained final composite models using all available datasets for each treatment type and these are ready to be applied to any additional dataset, such as the TBCRC 030^25^ (NCT01982448) and RESPONSE (NCT05020860) clinical trials.

From a clinical-utility standpoint, the cutpoint analyses in **Table 2** illustrate two distinct use cases. A high-specificity, rule-in threshold which is exemplified by the protein-informed platinum + taxane predictor, which flagged pCR tumors with 100% positive predictive value. This could identify patients likely to achieve pathologic complete response with chemotherapy alone, supporting de-escalation by sparing additional agents such as pembrolizumab and anthracycline/cyclophosphamide. Conversely, a high-sensitivity screening threshold, as applied to the taxane predictor in BEAUTY, prioritizes capturing as many responders as possible to minimize the risk of withholding an effective therapy. Realizing either application will require prospective validation in larger, uniformly annotated cohorts, but the ability to tune a single score toward ruling tumors in or out underscores the translational potential of these predictors.

Our previous WGCNA/CTD work employed pseudo-bulk RNA-Seq data in which the mouse stromal reads were added to the human epithelial tumor cell reads in an attempt to mimic bulk RNA-Seq data from human-only clinical samples. Stromal reads derived from murine stroma capture conserved fibroblast and vascular biology^33,34^, while human immune microenvironment features are not represented. Unfortunately, not all datasets were appropriate for such analysis. However, going forward, using stromal genes from pseudo-bulk data could also help improve our predictions as these genes have been demonstrated to be highly informative, especially for taxane response prediction, in our previous study that used a complementary approach (WGCNA/CTD) for chemotherapy response prediction^14^. Together with a recently described high-performance 20 gene biomarker panel to predict pCR to neoadjuvant taxane plus anthracycline-based chemotherapy regimens in TNBC^27^, these predictors cover the multitude of chemotherapy regimens given to TNBC patients and provides strong rationale for incorporation into future clinical trials. Models of negative biomarker combinations will also be very useful to predict progressive disease or residual cancer burden, but will require more studies that include these measurements together with molecular profiling.

Despite finding differences in molecular associations to various chemotherapy agents described above, our investigation into associations with response to any treatment arm tested identified KRT5, a basal cytokeratin associated with basal-like breast cancer, as a potential biomarker to stratify TNBC tumors for treatment. Immunohistochemistry for basal cytokeratins 5/6, 14, and 17 have been used previously to define basal-like breast cancer and examine associations with chemotherapy responses to different treatment regimens^35–38^. Further stratifying TNBC tumors using positivity for at least one of these basal cytokeratins identified patients with better adjuvant response to cyclophosphamide, methotrexate, and 5-fluorouracil (CMF) chemotherapy^36^ and to capecitabine^35^ but also identified patients with worse adjuvant response to anthracycline-based chemotherapy^37,38^ compared to patients without these markers. The presence of these markers was also associated with worse response of TNBC patients treated with anthracycline-based chemotherapy in the neoadjuvant setting^39^. Here, we observed high KRT5 protein abundance by mass-spectrometry-based measurements in TNBC tumors responsive to non-anthracycline-based chemotherapy consisting of single agent carboplatin, docetaxel, or their combination. These data suggest basal-like markers may be either positive or negative markers of TNBC response to chemotherapy depending on the regimen and also provide rationale for further investigation for KRT5 as a companion diagnostic. We confirmed KRT5 findings by IHC in PDX tumors and a KRT5 Allred IHC score achieved a high AUROC of 0.83 for discriminating responsive PDXs from this study to any chemotherapy (**Figure 3H**). Additional validation of KRT5 IHC in human breast cohorts with chemotherapy response information is ongoing. Future diagnostic assays and prospective clinical trials will be required to evaluate the clinical utility of these candidate biomarker or biomarker combinations.

In PDXs resistant to all treatment arms we also found elevated levels of OXPHOS and lower levels of glucose metabolism/glycolysis related pathways (**Figure 2D**). These findings of high OXPHOS but low glycolysis in our resistant PDXs is consistent with a previous study that also showed elevated OXPHOS and decreased glycolysis in a residual TNBC PDX tumor post adriamycin/cyclophosphamide chemotherapy treatment that was generated from a treatment-naive TNBC patient^40^. In addition, mitochondrial gene expression and protein synthesis were among the top pathways enriched in resistant PDXs, as was OXPHOS, which may be fueled by increased respiratory chain complex components (**Figure S3C**), consistent with previous reports that mitochondrial translation and oxidative phosphorylation support chemotherapy resistance and represent therapeutic vulnerabilities in TNBC^41,42^.

Although using PDXs in our study addresses many challenges in TNBC, it faces limitations that may impact clinical translation. Immune-deficient PDX models lack an effective immune response, a critical biological process that is well-established to become altered during cancer. The presence of specific immune cells in human tumors, such as tumor infiltrating lymphocytes (TILs), have been shown to be not only predictive of response to some chemotherapeutics, but also prognostic for TNBC patients^43,44^. This is particularly relevant to our KRT5-associated findings, as basal cytokeratin-high TNBCs in patients have been linked to higher tumor mutational and neoantigen burden and are enriched for TILs^45,46^, with TIL abundance itself serving as an independent predictor of chemoresponse and survival^47,48^. Another immunity-related issue is our previous observation where immunologically “cold” TNBC tumors have higher engraftment rates as PDX models compared to immune-enriched “hot” TNBC tumors^49^ leading to the vast majority of the TNBC models analyzed in the current study to be immunologically “cold”. It is not completely understood what the impact of an intact immune system would have on our study’s findings. However, we found groups of genes predictive of chemotherapy response (**Figures 4-5**). Similarities between PDX models and patients were also highlighted in our previous study that showed highly concordant drug responses between PDX and patient-of-origin, especially for taxanes^14^. In addition, our study investigated tumor responses after four cycles of chemotherapy similar to evaluating patient responses to neoadjuvant chemotherapy. Evaluation of longer-term measures such as tumor recurrence is an ongoing area of investigation by our team.

In conclusion, our proteogenomic characterization identifies candidate molecular mechanisms underlying response and resistance, as well as putative predictive biomarkers for stratifying TNBC tumors for single or combination chemotherapy treatments. These results also support evaluation of rationally selected PDX most likely to respond to such agents (or not) to a meaningful extent, and provide a valuable resource for researchers and clinicians.

## Methods

### RESOURCE AVAILABILITY

#### Lead Contact

Further information and requests should be directed to and will be fulfilled by the corresponding author, Michael. T. Lewis

### Materials Availability

Xenografts are available from the corresponding author for academic/nonprofit use on a cost recovery basis via a Material Transfer Agreement.

### Data availability statement

Genomic data have been deposited in the sequence read archive identifier PRJNA756268. RNA-Seq data have been deposited with the NCBI gene expression omnibus (GEO) accession number GSE183187. The mass spectrometry proteomics data have been deposited to the ProteomeXchange Consortium via the PRIDE^50^ partner repository with the dataset identifier PXD035857. Sample annotations, processed and normalized data files are provided in **Table S1B** and **Table S2**. Software and code used in this study are referenced in their corresponding Methods section. Publicly available and restricted TNBC datasets used for chemotherapy response prediction are described in **Table 1**. All other data are available upon reasonable request to corresponding author.

### Code availability statement

Final, locked chemotherapy-response predictor models (platinum, taxane, and platinum + taxane), together with code to apply them to new gene-expression data, are available at https://anonymous.4open.science/r/Lei-et-al---TNBC-chemo-predictors-1848. All other codes are available upon reasonable request to corresponding author.

## EXPERIMENTAL MODELS AND SUBJECT DETAILS

### Patient recruitment

Patients with breast cancer were recruited from clinics in the Lester and Sue Smith Breast Center at Baylor College of Medicine (BCM) (Houston, TX, USA) and Ben Taub General Hospital (Houston, TX, USA) under Institutional Review Board-approved protocols. Most patients received initial core-needle biopsies at the time of diagnosis and again either during or after treatment.

### Generation and chemotherapy response of PDX models

All human tissues were processed in compliance with NIH regulations and institutional guidelines, reviewed and approved by the Institutional Animal Care and Use Committee at Baylor College of Medicine. Single agent carboplatin, docetaxel, and untreated/vehicle control data were derived from three separate preclinical trials (**Figure S1B**) with further details as described previously in Petrosyan et al^14^. Response to chemotherapy was assessed by tumor volumes measured at pre- and post-treatment timepoints. Log_2_ transformed fold change in tumor volume at 28 days after initiation of treatment was calculated. If the tumor completely disappeared after treatment, a tumor volume of 0.05 was used in the calculation to avoid log transformation error. After log_2_ transformation, a general linear model (GLM) within each PDX was computed to compare the differences between treatment groups, and to estimate the average log_2_ fold change in tumor size and its associated 95% confidence interval for each treatment within a PDX. These data were also plotted for visual presentation. Response was classified based on modified RECIST 1.1 criteria^16,17^: Complete response (CR) was defined as non-palpable tumors. Partial response (PR) was defined as PDXs with ≥ 30% reduction in tumor volume but not reaching the non-palpable state. Stable disease (SD) was defined in PDXs with < 30% decrease but no more than 20% over baseline tumor volume. Progressive disease (PD) was defined for PDXs with tumor volume increases of > 20% over baseline tumor volume. Thresholds for quantitative responses were also determined. For example, since partial response (PR) is defined as ≥ 30% decrease in tumor volume below baseline at day 28 of treatment, a PDX model with an average log_2_ fold change in tumor size of < −0.515 [30% shrinkage ∼ log_2_(0.7)] would be deemed PR. mRECIST categories were assigned at the PDX level using the GLM-estimated mean log_2_ fold change as the point estimate. The associated 95% confidence intervals were reported for visualization but were not used for classification.

### Drug combination effect using Bliss independence model

To evaluate whether the carboplatin + docetaxel combination produced effects exceeding those expected under independent drug action, we re-analyzed average log_2_ fold-change (log_2_FC) estimates and 95% confidence intervals (CIs) derived from the within-PDX general linear models described above (the same estimates plotted in **Figures 1A and 1B**). Under Bliss independence^18^, unaffected fractions multiply (*U_AB_* = *U_A_* · *U_B_*), which on the log_2_FC scale is log-additivity relative to vehicle:

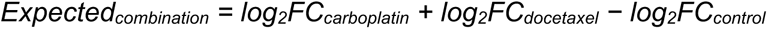

For each PDX with all four arms (control, carboplatin, docetaxel, and carboplatin + docetaxel; *n* = 42), we computed Expected and Observed combination log_2_FC values and tested the cohort-level difference using paired *t* and Wilcoxon signed-rank tests (**Figure S1C**). Individual PDX standard errors on the deviation *δ* = Observed − Expected were propagated from 95% CIs reported above as SE_arm_ = (Upper_CL − Lower_CL) / (2 × 1.96), with SE(*δ*) = √(SE_combination_^2^ + SE_carboplatin_^2^ + SE_docetaxel_^2^ + SE_control_^2^). CIs on *δ* were used to classify each PDX as more effective than predicted (95% CI entirely below zero), less effective than predicted (95% CI entirely above zero), or not significantly different from the prediction (95% CI crosses zero) (**Figure S1D**).

### Immunohistochemistry

PDX tumors were harvested, formalin-fixed for 24 hours, and then preserved in 70% ethanol. Tissue was paraffin embedded, cut into 3 µm sections, and immunohistochemical staining was performed. During IHC, antigen retrieval was performed using 0.1M citrate buffer pH 6.0 and sections were incubated with mouse anti-cytokeratin 5 monoclonal antibody (MA5-12596, Invitrogen) at 1:20 dilution for 1 hour at room temperature. Sections were incubated with HRP anti-mouse antibody (DAKO) and detected with DAB+ solution (DakoCytomation) and DAB sparkle enhancer (Biocare). Counterstain was performed using Harris modified hematoxylin. Images were acquired using a Nikon Eclipse Ci microscope with DS-Fi3 camera and NIS-Elements software. Staining for cytokeratin 5 was scored using the Allred method as previously described^51^.

## METHOD DETAILS

### Whole exome sequencing (WES)

DNA sequencing was performed by core facilities at BCM and Cornell University. Exome Sequencing Libraries were prepared using the Agilent SureSelect XT v6.0 Human kit. Sequencing was performed on an Illumina HiSeq 4000 machine (PE, 2x100 cycles, ∼133X estimated coverage per sample) at Cornell. At BCM, sequencing was performed on the Illumina NovaSeq machine (PE, 2x100, ∼200M read pairs per sample). Separation of human and mouse reads, alignment to the reference genome, variant calling, and annotation were performed using the PDXNet Tumor-Only Variant Calling pipeline developed by the Jackson Laboratory and hosted on the Cancer Genomics Cloud.

### RNA sequencing

Total RNA was extracted from PDX samples and 10 ng was used to generate and amplify transcriptome cDNA (NuGen Ovation v2, NuGen Technologies). Three grams of cNDA from each sample was fragmented to 250- 400 bases using the Covaris S2 focused ultrasonicator (Covaris). Using the Illumina TruSeq or NovaSeq DNA-Seq library preparation kits (Illumina Technologies), a double-stranded DNA library was generated with 1 g of the sheared cDNA. The library was quantified with a Kapa quantitative PCR Library Quantification Kit (Kapa). DNA libraries (11 pM) were loaded onto an Illumina HiSeq 2000 or NovaSeq6000 FlowCell and clusters were generated on the Illumina cBot. Paired-end 100-bp reads were generated on a HiSeq 2000 or NovaSeq6000 Sequencing System.

### Mass spectrometry methods

The tissues were pulverized in homemade kapton tube bags under liquid nitrogen to form powder and directly digested in 50 mM ammonium bicarbonate solution using trypsin enzyme at 37^0^C overnight. The digest was acidified with 10% formic acid (FA) and peptides were measured using the Pierce™ Quantitative Colorimetric Peptide Assay (Thermo Scientific 23275). 50 µg peptides were subjected to manual offline fractionation using a high pH reversed-phase chromatography to form a 5 fraction pool of peptides as described earlier^52^. LC-MS/MS analysis was carried out using a nano-LC 1200 system (Thermo Fisher Scientific, San Jose, CA) coupled to Orbitrap Fusion™ Lumos ETD mass spectrometer (Thermo Fisher Scientific, San Jose, CA). The fractionated peptides (1 µg) were loaded on a 2 cm X 100 µm I.D. trap-column (Reprosil-Pur Basic C18 1.9 µm beads, Dr.Maisch GmbH, Germany), eluted using a 75 min gradient of different concentration of buffer B (90% acetonitrile in 0.1% FA) as follows: [3% to 28%B - 70 min, 95%B - 5 min] at a flow rate of 750 nL/min on a 5 cm x 150 µm I.D. analytical column (Reprosil-Pur Basic C18 1.9 µm, Dr.Maisch GmbH, Germany). The peptides were directly electro-sprayed at 2.2 kV voltage into the mass spectrometer operated in a data-dependent mode with ‘top 30’ method. The full MS scan was acquired in Orbitrap in the range of 300-1400 m/z at 120,000 resolution (AGC 5E5, maxIT 50 ms) followed by MS2 in Ion Trap (HCD 28% collision energy, AGC 5E3, maxIT 35 ms) with 15 sec dynamic exclusion time.

## QUANTIFICATION AND STATISTICAL ANALYSIS

Statistical analyses were performed using R. Unless explicitly stated otherwise, all reported p-values are nominal (unadjusted).

### Genomic data analysis

#### Deleterious variant calling

All raw FASTQ files were subjected to QC verification by FASTQC (https://www.bioinformatics.babraham.ac.uk/projects/fastqc/) and were trimmed for adapter sequences with TrimGalore (https://www.bioinformatics.babraham.ac.uk/projects/trim_galore/). Whole Exome FASTQ files were processed using the Tumor-Only Variant Calling pipeline^53^ developed by the Jackson Laboratory for PDXNet hosted on the Cancer Genomics Cloud. This pipeline uses Xenome^54^ to separate human epithelial reads and mouse stromal reads, then aligns human reads against the GRCh38 human genome using BWA^55^. It then uses GATK MuTect2^56,57^ to call variants and SnpEff/SnpSift^58,59^ to annotate variants, generating per-sample VCF and tabular outputs for downstream analyses. Further annotation with gnomAD^60^ and CLINVAR^61^ was performed which added information by matching locus fields (Chromosome, Position, Reference and Alternate alleles) to standard annotation VCF files. Multi-allelic sites were decomposed to ensure Alternate alleles matched exactly with annotation sources.

#### Copy number calling

A set of 6 normal breast tissue samples were subjected to Whole Exome Sequencing both at Cornell and Baylor to serve as platform-matched normal samples. These platform-matched normal samples were run through the Variant Calling pipeline to obtain platform-matched normal BAMs. CopywriteR^62^ was used on each tumor BAM file obtained from the Variant Calling pipeline, and a random platform-matched normal BAM was used as control. CopywriteR uses depth information from off-target reads to calculate segment-level copy number data. This segment level data was then processed using GISTIC2.0^63^ to obtain raw and threshold gene-level and focal-level copy number information.

#### Mutational signature analysis

SBS mutational signatures were derived for each PDX from WES somatic variant calls (GRCh38/hg38). As the models lack matched normal tissue, somatic-like SNVs were approximated by retaining PASS-filtered single-nucleotide variants and removing likely germline polymorphisms (population allele frequency ≥ 0.1% in gnomAD exomes or 1000 Genomes Phase 3), yielding a median of 109 SNVs per model. Mutations were tallied into the 96 trinucleotide contexts and refit to COSMIC v3.4 SBS reference signatures (GRCh38) using MutationalPatterns (v3.14.0) with the BSgenome.Hsapiens.UCSC.hg38 reference. Refitting was restricted to a breast-carcinoma-relevant panel (SBS1, 2, 3, 5, 6, 8, 13, 17a, 17b, 18, 20, 26, 30) and performed with strict refitting (fit_to_signatures_strict, max_delta = 0.004) to limit over-assignment of the flat SBS3 signature in sparse exome spectra; contributions were expressed per model as each signature’s relative fraction. COSMIC SBS3 was used as the canonical marker of HRD. Robustness of SBS3 estimates was assessed over 100 bootstrap resamples of each spectrum (fit_to_signatures_bootstrapped, method = “strict”).

#### HRD association with response

Two composite HRD features were tested against carboplatin response using the same Fisher’s Exact Test framework as the per-gene deleterious-variant analysis (**Figure S2A**). Genetic HRD was defined as a deleterious variant in BRCA1 or BRCA2 (n = 16 PDX). The SBS3 signature was included as a binary feature by dichotomizing SBS3 relative contribution at the cohort median (“SBS3-high”, > 0.36; n = 25). For each feature, enrichment between responders (CR/PR) and non-responders (SD/PD) was tested by two-sided Fisher’s Exact Test on the 2×2 table of feature status × response, with the same orientation as the per-gene analysis (negative log₂ odds ratio = enrichment among responders). Genetic HRD was significantly enriched among carboplatin responders (log_2_ odds ratio = −2.9, p = 0.005), whereas SBS3-high status was not (log_2_ odds ratio = −0.2, p = 1.0). As a continuous sensitivity analysis, SBS3 relative contribution was also tested against day-28 log_2_ fold-change by Spearman’s correlation and was not associated with response to any regimen (carboplatin ρ = −0.03, p = 0.84; docetaxel ρ = −0.13, p = 0.37; combination ρ = −0.21, p = 0.19).

### RNA-Seq quantification

RNA-Seq FASTQ files were processed using Xenome to separate murine stromal reads and human epithelial reads, which were both then quantified with rsem-calculate-expression. A reference index was created for RSEM (v1.3.0)^64^ using rsem-prepare-reference on the hg38 and mm10 genome assembly (FASTA) and transcript feature (GTF) files to serve as library input to the rsem-calculate-expression tool. Expected counts from RSEM were collated to create a genes-by-samples matrix. This matrix was Upper-Quantile normalized per-sample to account for differences in sequencing depth and then log_2_ transformed to reduce expression variance across orders of magnitude.

### Proteomic data processing

Raw MS/MS data were processed using MaxQuant (version 1.6.5.0)^65^ with the Andromeda search engine3 against the human plus mouse RefSeq database. The standard MaxQuant contaminant database was also included in the searching. Enzyme specificity was set to trypsin and up to two missed cleavages were allowed. Oxidation (M), acetyl (protein N-term) and deamidation (NQ) were set as variable modifications. No fixed modification was selected. The maximum number of modifications per peptide was set as 4. The match between runs function was enabled. A false discovery rate cutoff of 1% was used at the PSM and protein levels. Reverse and contaminant matches were removed from the analysis. The file “evidence.txt” generated by MaxQuant was processed using gpGrouper^52^ to generate gene level quantification data. Different batch and normalization methods were evaluated using OmicsEV^66^ (https://github.com/bzhanglab/OmicsEV) which selected the following procedure to generate proteomic data for all downstream analyses: Gene products which were identified and quantified in at least two replicates of any PDX model were retained. Expression values were averaged across all replicates of each PDX model for which a measurement was made then batch correction was applied using ComBat from the sva R package^67^ across the cohort stratified by year of sample processing.

### Proteogenomic analysis

#### Publicly available datasets for molecular profiles of TNBC PDX and TNBC tumors

Molecular profiling data of baseline TNBC tumors prior to neoadjuvant chemotherapy was retrieved as follows. Microarray data for DFCI cohort (GSE18864)^68^ and ISPY2 (GSE194040)^69^ were downloaded from GEO, log_2_ transformed, and collapsed to gene level by mean probe expression. RNA-Seq data for TNBC PDXs from Savage et al (GSE142767)^23^ was also retrieved from GEO. RNA-Seq and Tandem Mass Tag (TMT) mass spectrometry-based proteomics data for CPTAC-TNBC trials was downloaded from supplemental tables in the Anurag et al, 2022 manuscript^10^. Controlled-access datasets were obtained directly from study investigators.

#### Molecular features and drug response associations

For associations with deleterious variants and copy number events, PDXs were binarized into responsive (CR + PR) and resistant (SD + PD) models according to each PDX’s mRECIST classification, assigned from the point estimate of the GLM-estimated average log_2_ fold change in tumor volume. Fisher’s exact test was performed to test for enrichment of deleterious variants in responsive or resistant models. Fisher’s exact test was also performed to test for enrichment of amplification (GISTIC = +2) or deep deletion (GISTIC = −2) events in responsive or resistant models. For both mutation and copy number data, Inf and –Inf values were set to 5 and –5, respectively. Spearman’s correlation was computed for genes quantified in at least 50% of PDXs by RNA-Seq and proteomics data with quantitative tumor measurements after treatment. For pathway analysis, signed −log_10_ p-values (positive sign if positively correlated with response, negative sign if negative correlated with response) from correlation analyses were used as input for gene set enrichment analysis (GSEA) with gene sets in Gene Ontology: Biological Processes, Reactome, and Wikipathway databases using WebGestalt^70^. A weighted set cover was then applied to reduce the number of genesets from each database to a minimal set while maximizing gene coverage^71^. Wilcoxon-rank sum and Anderson-Darling tests (using ad_test from the twosamples R package^72^) were performed for two group comparisons where indicated.

#### Molecular profiling association with drug response in external TNBC PDX and human TNBC datasets

For **Figure 2**, TNBC PDXs and human TNBC tumors from public datasets with available mRNA data and proteomic profiling were analyzed. Quantitative response metrics (estimated average log_2_ fold change or BestAvgResponse) or qualitative classifications responsive (CR/pCR or CR/PR) versus resistant (all else) were used for association with response to molecular profiles.

### Feature selection methods and generation of chemotherapy response predictors

Datasets used for training and evaluation of chemotherapy response predictors are listed in **Table 1**. Where necessary, datasets containing multiple breast cancer subtypes were restricted to TNBC samples prior to analysis. The Protein Marker Selection (ProMS) tool^22^ was initially used as a state-of-the-art feature selection method developed by our group to examine the performance of individual datasets to predict CR/pCR. ProMS was implemented in the Python package *proms* (https://pypi.org/project/proms) to select non-redundant and therefore complementary RNA markers to predict CR/pCR using mRNA alone. It identifies markers that represent distinct biological functions or pathways defined by co-expressed genes. TNBC samples from PDXs and clinical trial samples with mRNA were categorized to be responsive (CR or pCR) or resistant (all else). Performance of ProMS-selected markers on individual datasets was evaluated by training logistic regression classifiers under Monte-Carlo Cross Validation (MCCV) over 50 iterations. In each iteration, the dataset was randomly split 70% for classifier training (T) and 30% for validation (V). Within T, ProMS selected k = 5 mRNA markers and a logistic regression classifier was fit on those markers. A number of hyperparameters were tuned, including univariate filtering and L2 regularization (grid search: 0.001, 0.01, 0.1, 1, 10, and 100), by 3-fold cross-validation within T. All other parameters were used as default, the details of which can be found in the ProMS manuscript^22^.

Since models trained on individual datasets were unstable and not highly predictive, with Dataset 07^23^ (**Table 1**) showing particularly poor performance (AUROC = 0.35) and therefore excluded from subsequent analyses, datasets of the same treatment type (platinum: carboplatin and cisplatin; taxane: docetaxel and paclitaxel; platinum + taxane: carboplatin + docetaxel and carboplatin + paclitaxel)use were batch corrected using ComBat^67^ and merged to increase sample size. Four statistically diverse feature selection methods were then employed on the merged datasets with the test dataset held out (**Table 1**, Role: Figure 4 column) to identify transcriptomic biomarkers predictive of chemotherapy response.

#### Method 1: Regression genes

Genes whose mRNA expression levels were significantly associated with tumor response were identified from merged datasets. For datasets where quantitative tumor response measurements were available, Spearman’s correlation between gene expression and estimated log_2_ fold change in tumor volume was used (p < 0.05). For datasets where only qualitative response classifications were available, genes significantly differentially expressed between responsive and resistant groups were used instead (p < 0.05). Genes reaching significance in a minimum number of training datasets were retained: ≥3 datasets for platinum, ≥2 for taxane, and ≥2 for combination predictors. For the platinum predictor, genes previously implicated in platinum resistance in cancer from xenograft or patient tissues^19^ were additionally included as a training dataset.

#### Method 2: WGCNA-CTD

Weighted gene co-expression network analysis followed by Connect The Dots (WGCNA/CTD) was used to identify mRNA-based biomarker panels as previously described by our group^14^. This approach groups genes into co-expression modules and then applies CTD to identify genes within those modules that are most informative for distinguishing responsive from resistant tumors.

#### Method 3: ProMS

The same ProMS procedure described above was applied to the merged, batch-corrected datasets to select *k* = 5-10 markers to predict CR/pCR and CR/PR, evaluated by MCCV 25 times. The number of markers giving the highest average MCCV AUROC was selected to repeat the feature selection and classifier building process using all data to produce a final fully trained, fixed model whose performance was assessed by AUROC on independent test data. In multi-omics mode (ProMS^mo^), protein data were incorporated alongside RNA data to guide RNA marker selection, enabling protein-informed feature identification. For the platinum + taxane predictor, proteins quantified in all samples were used to generate ComBat batch-corrected merged protein data and used together with merged mRNA data to select 10 protein-informed, mRNA markers.

#### Method 4: CR/PR- or (p)CR-associated genes

Genes whose mRNA expression was significantly different between responsive and resistant groups were identified by differential expression analysis. For objective response (CR/PR) predictors, genes differentially expressed between CR/PR and SD/PD groups were selected. For pathologic/complete response features, genes differentially expressed between CR/pCR and all other response categories were selected. A significance threshold of p < 0.05 was applied. Genes reaching significance in a minimum number of training datasets were retained: ≥3 datasets for platinum, ≥2 for taxane, and ≥2 for combination predictors.

Each of the four methods generated feature sets which were subsequently used for classifier development and evaluation. For regression genes, WGCNA-CTD, and CR/PR-associated genes, logistic regression classifiers were trained using these features with MCCV (50 iterations, 70% training [T] / 30% validation [V] stratified split). In each iteration, L2 regularization was tuned by 3-fold cross-validation within T over the grid 0.001, 0.01, 0.1, 1, 10, 100, and AUROC was evaluated on V. The average AUROC across the 50 iterations was reported as the MCCV performance estimate. The most frequent best-fitting L2 value across the 50 iterations was then used to refit logistic regression on all training samples, producing the locked model. For ProMS, performance was estimated during the k-sweep MCCV described in Method 3. The locked model was produced by running ProMS once on all training samples at the selected k, with logistic regression fit on those features and L2 regularization tuned by 3-fold cross-validation over the same grid. Features from all four methods were also pooled into a combined feature set per treatment type, and a logistic regression classifier was trained on the pooled set using the same MCCV procedure described above for the three non-ProMS methods. The locked models were applied to the independent test set (the largest clinical dataset for that treatment type was withheld for this purpose, **Table 1**, Role: Figure 4 column) to evaluate fully trained model performance. To generate final composite models using as much data as possible, previously withheld clinical datasets were incorporated into training (**Table 1**, Role: Figure 5 column). Logistic regression classifiers were then trained using the same MCCV procedure described above, separately for each of the four feature sets and for a pooled feature set combining all four. Internal performance was estimated from the held-out validation set in each iteration. Performance of these fully trained composite predictors was assessed on datasets that were never used at any training stage throughout the entire study: the clinical BEAUTY cohort for the taxane predictor and an additional cohort of 11 PDXs for the platinum and platinum + taxane predictors. The BEAUTY cohort and the 11-PDX cohort from **Table 1** were not included in the ComBat batch correction of the training cohorts. Each was independently standardized before the fixed, composite models were applied.

### Classification cutpoints

To derive candidate clinical decision rules, predicted probabilities from the locked models on each independent test set were converted to binary calls using two complementary strategies (**Table 2**). The screening cutpoint was defined as the highest sensitivity attainable away from the tail of the ROC curve and, among the candidate cutpoints achieving that sensitivity, the one with the highest specificity, favoring detection of responders (i.e., ruling out non-responders). The diagnostic cutpoint was defined as the score at which sensitivity and specificity, each plotted as a function of the cutpoint, intersect, thereby maximizing the joint (sensitivity, specificity) pair. At each cutpoint, samples with a predicted probability greater than the threshold were classified as predicted responders, and sensitivity, specificity, and positive predictive value were computed against the observed response labels. Exact Clopper-Pearson binomial 95% confidence intervals were reported for sensitivity and specificity. These cutpoints were derived post hoc on the independent test data and are intended as illustrative operating points requiring prospective validation.

## Acknowledgments

We honor the breast cancer patients who donate their biopsies for PDX generation. Ms Susan Rafte provided research advocacy support for this work. This work was supported by NIH/NCI grants U54 CA224076 (to M.T.L, A.L.W., and B.E.W.), U24 CA226110 (to M.T.L.), U24 CA210954 and U24 CA271076 (both to B.Z.), R37 CA269783-01A1 (to G.V.E.), K22 Career Transition Award 1K22CA241113-01 (to G.V.E.), SPORE P50 CA186784, Cancer Center Support Grant P30 CA125123, by a Translational Breast Cancer Research Training Program grant T32 CA203690 (to J.T.L.), and by Core Facility grants from the Cancer Prevention and Research Institute of Texas (CPRIT) RP170691 and RP220646 (both to M.T.L.), NIH 1S10OD023469 and CPRIT RP200504 (both to D.C.K.), and additionally by a CPRIT Recruitment of Rising Stars Award RR160027 (to B.Z.) and CPRIT First time-faculty recruitment grant RR200009 (to G.V.E.), and an American Cancer Society Research Scholar Grant RSG-22-093-01-CCB (to G.V.E.). N.Z. was supported by CPRIT RP220468. M.A. acknowledges support from U01CA214125 and NCI-SPORE P50 CA186784-06. J.C.B. is supported by the W.H. Odell Professorship in Individualized Medicine. This work was also supported by a grant from the V Foundation and a generous gift from the Korell family for the study of triple-negative breast cancer, the Mayo Clinic Center for Individualized Medicine, the Nadia’s Gift Foundation, John P. Guider, the George M. Eisenberg Foundation for Charities, and the Eveleigh Family (to M.P.G., L.W., and J.C.B.). Additional support was provided by the Mayo Clinic Cancer Center (CA15083-40A2) and the Mayo Clinic Breast Specialized Program of Research Excellence (SPORE P50 CA116201, to M.P.G., V.J.S., K.R.K., and L.W.). The BCM Mass Spectrometry Proteomics Core is supported by the Dan L Duncan Comprehensive Cancer Center NIH award P30 CA125123 and a CPRIT Core Facility Award RP17005. M.T.L, B.Z., and G.V.E. are CPRIT Scholars in Cancer Research, and B.Z. is also a McNair Scholar. M.P.G. is the Erivan K. Haub Family Professor of Cancer Research Honoring Richard F. Emslander, M.D.

## Author Contributions

Conceptualization, M.T.L., B.Z., G.V.E., M.A., and J.T.L.; Data collection and processing, L.E.D., A.N.L., A.B.S., A.M., C.S., N.Z., J.C., R.R.S., D.C.K., B.W., C.H., A.J., K.L., J.B.P., and J.T.L.; Data analysis, J.T.L., M.T.L., B.Z., R.R.S., C.H., S.V.V., T.W., C.I.M., K.R.K., V.J.S., Y.L., and V.P.; Data curation, R.R.S.; Validation, L.E.D. and J.D.L.; Resources, M.T.L., B.Z., S.L., M.J.E., M.F.R., S.G.H., G.M.W., E.M., S.D., X.Z., F.M-B., X.C., M.P.G., J.C.B., L.W., B.E.W., and A.L.W.; Writing - original draft: J.T.L., M.T.L., and B.Z.; Funding acquisition: M.T.L., A.L.W., B.E.W., and B.Z.; Supervision, M.T.L. and B.Z.

## Supplemental Figure Legends

**Figure S1. Combination carboplatin and docetaxel is largely ineffective at generating enhanced responses over the best single agent in TNBC PDXs. Related to Figure 1. A)** Overall study design. **B)** Three preclinical trials from which the 50 TNBC PDXs models in the “chemotherapy response cohort” were chosen. **C)** Boxplots of computed expected vs observed combination PDX tumor responses at day 28. Expected response computed under the Bliss independence model (**Methods**). Lines connect paired values from the same PDX. P- values from paired t-test and Wilcoxon signed-rank test. **D)** Per-PDX deviation δ between observed and Bliss- expected combination response at day 28. Each point is one PDX and error bars represent 95% CIs. Blue, combination more effective than predicted (p < 0.05); grey, not significantly different from prediction; red, combination less effective than predicted (p < 0.05). Dashed line at δ = 0 indicates a perfect match to the prediction.

**Figure S2. Molecular associates of carboplatin, docetaxel, and combination treatment. Related to Figure 2. A-C)** Association of individual omic platforms with tumor responses to carboplatin (A), docetaxel (B), and combination (C). With WES data, plots were generated showing significant enrichment (p < 0.05) of deleterious variants and high-level amplification events (GISTIC = +2) associated with responsive (blue dots) or resistant (red dots) PDXs, with p-values and odds ratios derived from Fisher’s Exact Test (dashed horizontal line indicates p = 0.05). No deep deletion events (GISTIC = –2) were found to be significantly associated with any treatment arm and therefore not shown. To evaluate homologous-recombination deficiency (HRD) directly, two composite HRD features (purple dots) were tested by the same Fisher’s Exact Test and overlaid: “genetic HRD”, defined as a deleterious variant in BRCA1 or BRCA2, and “SBS3-high”, defined by dichotomizing the COSMIC SBS3 (HRD) mutational-signature contribution at the cohort median. Genetic HRD was significantly enriched among carboplatin responders, whereas the SBS3 signature was not associated with response. With RNA-Seq and proteomics data, plots were generated showing significantly associated genes (p < 0.05) that are higher in PDXs with increasing responsiveness to treatment (blue dots) or higher in PDXs with increasing resistance to treatment (red dots), with p-values derived from Spearman’s correlation between RNA/protein measurements and quantitative log_2_ fold change in tumor responses.

**Figure S3. Associations with resistance to all chemotherapy arms. Related to Figure 2. A)** Association of individual omic platforms with PDXs that were resistant to all treatment arms (mean log_2_ fold change or 95% CI in SD or PD range) vs those that had response to any treatment arm (PR or better). PDX group stratification shown in **Figure 1A** for “Q2”. With WES data, plots were generated showing significant enrichment (p < 0.05) of deleterious variants and high-level amplification events (GISTIC = +2) associated with PDXs that had response to any treatment arm (blue dots) or PDXs resistant to all treatment arms (red dots) with p-values and odds ratios derived from Fisher’s Exact Test (dashed horizontal line indicates p = 0.05). No deep deletion events (GISTIC = –2) were found to be significantly associated with any treatment arm and therefore not shown. With RNA-Seq and proteomics data, plots were generated showing significantly associated genes (p < 0.05) higher in PDXs with a response to any treatment arm (blue dots) or higher in PDXs resistant to all treatment arms (red dots) with p-values derived from Wilcoxon rank-sum tests. **B-E)** Top 10 leading genes for fatty acid (B), oxphos (C), stress response (D), and glucose pathways (E).

**Figure S4. Predicting chemotherapy response. Related to Figure 4. A)** Training and evaluation strategy for chemotherapy response predictors. **B)** Area under ROC curve (AUROC) performance in the set-aside cross- validation (CV) data from the same cohort of trained logistic regression models repeated using ProMS feature selection (5 features) for predicting complete response/pathologic complete response. Numbers indicate average AUROC from 50 times of Monte-Carlo cross-validations. Datasets used are summarized in **Table 1**. **C- F)** PCA plots depicting platinum (C), taxane (D), and platinum + taxane combination (F) datasets with RNA and protein (F) data before and after merging datasets post batch correction with ComBat.

**Figure S5. Single- and multi-omic predictors of chemotherapy response. Related to Figure 4. A,D,G)** Area under the receiver operating curve (AUROC) performance in the set-aside cross-validation data repeated 25 times of trained logistic regression models using ProMS feature selection (5-10 features) for predicting complete response (CR)/pathologic complete response (pCR) or complete response plus partial response (CR/PR) as indicated from combined RNA datasets for single-agent platinum (A), single-agent taxane (D), and platinum + taxane combination (G). Numbers indicate average AUROC. The number of selected markers giving the highest average AUROC for the CR/pCR (solid) or CR/PR (dashed) predictors were used to generate final, fully trained models. Datasets used are summarized in **Table 1**. **B,E,H)** ProMS features selected features giving the highest average cross-validation AUROC from (A), (D), and (G). A multi-omics, protein-facilitated RNA marker selection method (ProMS^mo^) was also examined in (H) with 10 features which was based on the number of markers with the highest cross-validation AUROC in (G). **C,F,I)** Performance of fully trained logistic regression (LR) models using markers selected by ProMS/ProMS^mo^ in (B,E,H), respectively to predict chemoresponse in independent test data.

**Figure S6. Pathway enrichment of the chemotherapy-response predictor-gene sets. Related to Figures 4 and 5**. Pooled predictor genes (union of four feature-selection methods: regression, ProMS, WGCNA-CTD, and (p)CR-associated) used as predictive features shown in **Figures 4A, 4C, and 5A** were tested by over- representation analysis (ORA) for KEGG and Reactome gene sets in each treatment context (platinum, taxane, combination), and significant terms (FDR < 0.05) were reduced by weighted set cover. **A)** Selected pathways (rows) across the eight predictor-gene panels (columns; grouped by figure, labeled by treatment). Color = ORA FDR; asterisk = FDR < 0.05; row bar = database. **B)** Membership of predictor genes (rows) in the Panel A pathways (columns; black = gene in pathway). Left tracks show which feature-selection method(s) selected each gene (black = yes; not mutually exclusive); top bars give figure, treatment, and database.

## Supplemental Table Titles and Legends

**Supplementary Table S1**. Breast PDX enrollment from three preclinical trials and chemotherapy response and metadata, related to Figures 1 and S1.

**Supplementary Table S2**. Processed genomic, transcriptomic, and proteomic data associated with this study, related to Figures 2 and S2.

**Supplementary Table S3**. Association results for mutation, copy number, RNA-Seq, and proteomics with single agent and combination treatment, related to Figures 2 and S2.

**Supplementary Table S4**. Gene and pathway associations for resistant vs responsive PDXs, related to Figures 2 and S3.

**Supplementary Table S5**. mRNA genes selected by different feature selection methods for chemotherapy CR/PR response prediction, related to Figures 4A-B.

**Supplementary Table S6.** mRNA genes selected by different feature selection methods for chemotherapy (p)CR response prediction, related to Figures 4C-D.

**Supplementary Table S7.** mRNA genes selected by different feature selection methods for composite chemotherapy (p)CR response prediction using all possible datasets, related to Figures 5A-B.

## References

1. Engel, C., Rhiem, K., Hahnen, E., Loibl, S., Weber, K.E., Seiler, S., Zachariae, S., Hauke, J., Wappenschmidt, B., Waha, A., et al. (2018). Prevalence of pathogenic BRCA1/2 germline mutations among 802 women with unilateral triple-negative breast cancer without family cancer history. BMC Cancer 18, 265.

2. Bear, H.D., Anderson, S., Brown, A., Smith, R., Mamounas, E.P., Fisher, B., Margolese, R., Theoret, H., Soran, A., Wickerham, D.L., et al. (2003). The effect on tumor response of adding sequential preoperative docetaxel to preoperative doxorubicin and cyclophosphamide: preliminary results from National Surgical Adjuvant Breast and Bowel Project Protocol B-27. J. Clin. Oncol. 21, 4165–4174.

3. Mamounas, E.P., Bryant, J., Lembersky, B., Fehrenbacher, L., Sedlacek, S.M., Fisher, B., Wickerham, D.L., Yothers, G., Soran, A., and Wolmark, N. (2005). Paclitaxel after doxorubicin plus cyclophosphamide as adjuvant chemotherapy for node-positive breast cancer: results from NSABP B-28. J. Clin. Oncol. 23, 3686–3696.

4. Lehmann, B.D., Bauer, J.A., Chen, X., Sanders, M.E., Chakravarthy, A.B., Shyr, Y., and Pietenpol, J.A. (2011). Identification of human triple-negative breast cancer subtypes and preclinical models for selection of targeted therapies. J. Clin. Invest. 121, 2750–2767.

5. von Minckwitz, G., Schneeweiss, A., Loibl, S., Salat, C., Denkert, C., Rezai, M., Blohmer, J.U., Jackisch, C., Paepke, S., Gerber, B., et al. (2014). Neoadjuvant carboplatin in patients with triple-negative and HER2-positive early breast cancer (GeparSixto; GBG 66): a randomised phase 2 trial. Lancet Oncol. 15, 747–756.

6. Loibl, S., O’Shaughnessy, J., Untch, M., Sikov, W.M., Rugo, H.S., McKee, M.D., Huober, J., Golshan, M., von Minckwitz, G., Maag, D., et al. (2018). Addition of the PARP inhibitor veliparib plus carboplatin or carboplatin alone to standard neoadjuvant chemotherapy in triple-negative breast cancer (BrighTNess): a randomised, phase 3 trial. Lancet Oncol. 19, 497–509.

7. Sikov, W.M., Berry, D.A., Perou, C.M., Singh, B., Cirrincione, C.T., Tolaney, S.M., Kuzma, C.S., Pluard, T.J., Somlo, G., Port, E.R., et al. (2015). Impact of the addition of carboplatin and/or bevacizumab to neoadjuvant once-per-week paclitaxel followed by dose-dense doxorubicin and cyclophosphamide on pathologic complete response rates in stage II to III triple-negative breast cancer: CALGB 40603 (Alliance). J. Clin. Oncol. 33, 13–21.

8. Zheng, R., Han, S., Duan, C., Chen, K., You, Z., Jia, J., Lin, S., Liang, L., Liu, A., Long, H., et al. (2015). Role of taxane and anthracycline combination regimens in the management of advanced breast cancer: a meta-analysis of randomized trials. Medicine 94, e803.

9. Zhang, L., Wu, Z.-Y., Li, J., Lin, Y., Liu, Z., Cao, Y., Zhang, G., Gao, H.-F., Yang, M., Yang, C.-Q., et al. (2022). Neoadjuvant docetaxel plus carboplatin vs epirubicin plus cyclophosphamide followed by docetaxel in triple-negative, early-stage breast cancer (NeoCART): Results from a multicenter, randomized controlled, open-label phase II trial. Int. J. Cancer 150, 654–662.

10. Anurag, M., Jaehnig, E.J., Krug, K., Lei, J.T., Bergstrom, E.J., Kim, B.-J., Vashist, T.D., Huynh, A.M.T., Dou, Y., Gou, X., et al. (2022). Proteogenomic Markers of Chemotherapy Resistance and Response in Triple-Negative Breast Cancer. Cancer Discov. 12, 2586–2605.

11. Symmans, W.F., Wei, C., Gould, R., Yu, X., Zhang, Y., Liu, M., Walls, A., Bousamra, A., Ramineni, M., Sinn, B., et al. (2017). Long-Term Prognostic Risk After Neoadjuvant Chemotherapy Associated With Residual Cancer Burden and Breast Cancer Subtype. J. Clin. Oncol. 35, 1049–1060.

12. Schmid, P., Cortes, J., Pusztai, L., McArthur, H., Kümmel, S., Bergh, J., Denkert, C., Park, Y.H., Hui, R., Harbeck, N., et al. (2020). Pembrolizumab for early triple-negative breast cancer. N. Engl. J. Med. 382, 810–821.

13. Pusztai, L., Denkert, C., O’Shaughnessy, J., Cortes, J., Dent, R., McArthur, H., Kümmel, S., Bergh, J., Park, Y.H., Hui, R., et al. (2024). Event-free survival by residual cancer burden with pembrolizumab in early-stage TNBC: exploratory analysis from KEYNOTE-522. Ann. Oncol. 35, 429–436.

14. Petrosyan, V., Dobrolecki, L.E., Thistlethwaite, L., Lewis, A.N., Sallas, C., Srinivasan, R.R., Lei, J.T., Kovacevic, V., Obradovic, P., Ellis, M.J., et al. (2023). Identifying biomarkers of differential chemotherapy response in TNBC patient-derived xenografts with a CTD/WGCNA approach. iScience 26, 105799.

15. Kent, D.M., and Hayward, R.A. (2007). Limitations of applying summary results of clinical trials to individual patients: the need for risk stratification. JAMA 298, 1209–1212.

16. Zhang, X., Claerhout, S., Prat, A., Dobrolecki, L.E., Petrovic, I., Lai, Q., Landis, M.D., Wiechmann, L., Schiff, R., Giuliano, M., et al. (2013). A renewable tissue resource of phenotypically stable, biologically and ethnically diverse, patient-derived human breast cancer xenograft models. Cancer Res. 73, 4885–4897.

17. Meric-Bernstam, F., Lloyd, M.W., Koc, S., Evrard, Y.A., McShane, L.M., Lewis, M.T., Evans, K.W., Li, D., Rubinstein, L., Welm, A., et al. (2024). Assessment of patient-derived xenograft growth and antitumor activity: The NCI PDXNet consensus recommendations. Mol. Cancer Ther. 23, 924–938.

18. Bliss, C.I. (1939). The toxicity of poisons applied jointly. Ann. Appl. Biol. 26, 585–615.

19. Huang, D., Savage, S.R., Calinawan, A.P., Lin, C., Zhang, B., Wang, P., Starr, T.K., Birrer, M.J., and Paulovich, A.G. (2021). A highly annotated database of genes associated with platinum resistance in cancer. Oncogene 40, 6395–6405.

20. Savage, S.R., Yi, X., Lei, J.T., Wen, B., Zhao, H., Liao, Y., Jaehnig, E.J., Somes, L.K., Shafer, P.W., Lee, T.D., et al. (2024). Pan-cancer proteogenomics expands the landscape of therapeutic targets. Cell. 10.1016/j.cell.2024.05.039.

21. Park, J.H., Vithayathil, S., Kumar, S., Sung, P.-L., Dobrolecki, L.E., Putluri, V., Bhat, V.B., Bhowmik, S.K., Gupta, V., Arora, K., et al. (2016). Fatty acid oxidation-driven Src links mitochondrial energy reprogramming and oncogenic properties in triple-negative breast cancer. Cell Rep. 14, 2154–2165.

22. Shi, Z., Wen, B., Gao, Q., and Zhang, B. (2021). Feature Selection Methods for Protein Biomarker Discovery from Proteomics or Multiomics Data. Mol. Cell. Proteomics 20, 100083.

23. Savage, P., Pacis, A., Kuasne, H., Liu, L., Lai, D., Wan, A., Dankner, M., Martinez, C., Muñoz-Ramos, V., Pilon, V., et al. (2020). Chemogenomic profiling of breast cancer patient-derived xenografts reveals targetable vulnerabilities for difficult-to-treat tumors. Commun Biol 3, 310.

24. Goetz, M.P., Kalari, K.R., Suman, V.J., Moyer, A.M., Yu, J., Visscher, D.W., Dockter, T.J., Vedell, P.T., Sinnwell, J.P., Tang, X., et al. (2017). Tumor sequencing and patient-derived xenografts in the neoadjuvant treatment of breast cancer. J. Natl. Cancer Inst. 109. 10.1093/jnci/djw306.

25. Mayer, E.L., Abramson, V., Jankowitz, R., Falkson, C., Marcom, P.K., Traina, T., Carey, L., Rimawi, M., Specht, J., Miller, K., et al. (2020). TBCRC 030: a phase II study of preoperative cisplatin versus paclitaxel in triple-negative breast cancer: evaluating the homologous recombination deficiency (HRD) biomarker. Ann. Oncol. 31, 1518–1525.

26. Edlund, K., Madjar, K., Lebrecht, A., Aktas, B., Pilch, H., Hoffmann, G., Hofmann, M., Kolberg, H.-C., Boehm, D., Battista, M., et al. (2021). Gene expression-based prediction of neoadjuvant chemotherapy response in early breast cancer: Results of the prospective multicenter EXPRESSION trial. Clin. Cancer Res. 27, 2148–2158.

27. Inayatullah, M., Mahesh, A., Turnbull, A.K., Dixon, J.M., Natrajan, R., and Tiwari, V.K. (2024). Basal-epithelial subpopulations underlie and predict chemotherapy resistance in triple-negative breast cancer. EMBO Mol. Med. 16, 823–853.

28. Sharma, P., López-Tarruella, S., García-Saenz, J.A., Khan, Q.J., Gómez, H.L., Prat, A., Moreno, F., Jerez-Gilarranz, Y., Barnadas, A., Picornell, A.C., et al. (2018). Pathological response and survival in triple-negative breast cancer following neoadjuvant carboplatin plus docetaxel. Clin. Cancer Res. 24, 5820–5829.

29. Menghi, F., Banda, K., Kumar, P., Straub, R., Dobrolecki, L., Rodriguez, I.V., Yost, S.E., Chandok, H., Radke, M.R., Somlo, G., et al. (2022). Genomic and epigenomic BRCA alterations predict adaptive resistance and response to platinum-based therapy in patients with triple-negative breast and ovarian carcinomas. Sci. Transl. Med. 14, eabn1926.

30. Sharma, P., López-Tarruella, S., García-Saenz, J.A., Ward, C., Connor, C.S., Gómez, H.L., Prat, A., Moreno, F., Jerez-Gilarranz, Y., Barnadas, A., et al. (2017). Efficacy of neoadjuvant carboplatin plus docetaxel in triple-negative breast cancer: Combined analysis of two cohorts. Clin. Cancer Res. 23, 649– 657.

31. Burstein, H.J. (2024). Immunotherapy for early-stage triple-negative breast cancer. N. Engl. J. Med. 391, 2048–2049.

32. Kok, M., Gielen, R.-J., Adams, S., Lennerz, J.K., Sharma, P., Loibl, S., Reardon, E., Sonke, G., Linn, S., Delaloge, S., et al. (2024). Academic uphill battle to personalize treatment for patients with stage II/III triple-negative breast cancer. J. Clin. Oncol. 42, 3523–3529.

33. Isella, C., Terrasi, A., Bellomo, S.E., Petti, C., Galatola, G., Muratore, A., Mellano, A., Senetta, R., Cassenti, A., Sonetto, C., et al. (2015). Stromal contribution to the colorectal cancer transcriptome. Nat. Genet. 47, 312–319.

34. Kluin, R.J.C., Kemper, K., Kuilman, T., de Ruiter, J.R., Iyer, V., Forment, J.V., Cornelissen-Steijger, P., de Rink, I., Ter Brugge, P., Song, J.-Y., et al. (2018). XenofilteR: computational deconvolution of mouse and human reads in tumor xenograft sequence data. BMC Bioinformatics 19, 366.

35. Zhu, Y., Li, K., Zhang, J., Wang, L., Sheng, L., and Yan, L. (2021). The prognostic and predictive significance of cytokeratin 5/6 and epidermal growth factor receptor in metastatic triple-negative breast cancer treated with maintenance capecitabine. Transl. Cancer Res. 10, 1193–1203.

36. Rakha, E.A., Elsheikh, S.E., Aleskandarany, M.A., Habashi, H.O., Green, A.R., Powe, D.G., El-Sayed, M.E., Benhasouna, A., Brunet, J.-S., Akslen, L.A., et al. (2009). Triple-negative breast cancer: distinguishing between basal and nonbasal subtypes. Clin. Cancer Res. 15, 2302–2310.

37. Conforti, R., Boulet, T., Tomasic, G., Taranchon, E., Arriagada, R., Spielmann, M., Ducourtieux, M., Soria, J.C., Tursz, T., Delaloge, S., et al. (2007). Breast cancer molecular subclassification and estrogen receptor expression to predict efficacy of adjuvant anthracyclines-based chemotherapy: a biomarker study from two randomized trials. Ann. Oncol. 18, 1477–1483.

38. Cheang, M.C.U., Voduc, D., Bajdik, C., Leung, S., McKinney, S., Chia, S.K., Perou, C.M., and Nielsen, T.O. (2008). Basal-like breast cancer defined by five biomarkers has superior prognostic value than triple-negative phenotype. Clin. Cancer Res. 14, 1368–1376.

39. Abdelrahman, A.E., Rashed, H.E., Abdelgawad, M., and Abdelhamid, M.I. (2017). Prognostic impact of EGFR and cytokeratin 5/6 immunohistochemical expression in triple-negative breast cancer. Ann. Diagn. Pathol. 28, 43–53.

40. Echeverria, G.V., Ge, Z., Seth, S., Zhang, X., Jeter-Jones, S., Zhou, X., Cai, S., Tu, Y., McCoy, A., Peoples, M., et al. (2019). Resistance to neoadjuvant chemotherapy in triple-negative breast cancer mediated by a reversible drug-tolerant state. Sci. Transl. Med. 11. 10.1126/scitranslmed.aav0936.

41. Lamb, R., Ozsvari, B., Lisanti, C.L., Tanowitz, H.B., Howell, A., Martinez-Outschoorn, U.E., Sotgia, F., and Lisanti, M.P. (2015). Antibiotics that target mitochondria effectively eradicate cancer stem cells, across multiple tumor types: treating cancer like an infectious disease. Oncotarget 6, 4569–4584.

42. Berner, M.J., Wall, S.W., Baek, M.L., Lane, A., Greer, A.S., Wang, K., Dobrolecki, L.E., Strope, I., Zhu, Q., Zhang, B., et al. (2026). Mitochondrial translation is a targetable dependency of chemo-refractory triple negative breast cancer. bioRxivorg. 10.64898/2026.03.23.712647.

43. Loi, S., Sirtaine, N., Piette, F., Salgado, R., Viale, G., Van Eenoo, F., Rouas, G., Francis, P., Crown, J.P.A., Hitre, E., et al. (2013). Prognostic and predictive value of tumor-infiltrating lymphocytes in a phase III randomized adjuvant breast cancer trial in node-positive breast cancer comparing the addition of docetaxel to doxorubicin with doxorubicin-based chemotherapy: BIG 02-98. J. Clin. Oncol. 31, 860–867.

44. Wang, K., Xu, J., Zhang, T., and Xue, D. (2016). Tumor-infiltrating lymphocytes in breast cancer predict the response to chemotherapy and survival outcome: A meta-analysis. Oncotarget 7, 44288–44298.

45. Karn, T., Jiang, T., Hatzis, C., Sänger, N., El-Balat, A., Rody, A., Holtrich, U., Becker, S., Bianchini, G., and Pusztai, L. (2017). Association between genomic metrics and immune infiltration in triple-negative breast cancer. JAMA Oncol. 3, 1707–1711.

46. Karn, T., Denkert, C., Weber, K.E., Holtrich, U., Hanusch, C., Sinn, B.V., Higgs, B.W., Jank, P., Sinn, H.P., Huober, J., et al. (2020). Tumor mutational burden and immune infiltration as independent predictors of response to neoadjuvant immune checkpoint inhibition in early TNBC in GeparNuevo. Ann. Oncol. 31, 1216–1222.

47. Denkert, C., von Minckwitz, G., Darb-Esfahani, S., Lederer, B., Heppner, B.I., Weber, K.E., Budczies, J., Huober, J., Klauschen, F., Furlanetto, J., et al. (2017). Tumour-infiltrating lymphocytes and prognosis in different subtypes of breast cancer: a pooled analysis of 3771 patients treated with neoadjuvant therapy. Lancet Oncol. 19, 40–50.

48. Loi, S., Drubay, D., Adams, S., Pruneri, G., Francis, P.A., Lacroix-Triki, M., Joensuu, H., Dieci, M.V., Badve, S., Demaria, S., et al. (2019). Tumor-infiltrating lymphocytes and prognosis: A pooled individual patient analysis of early-stage triple-negative breast cancers. J. Clin. Oncol. 37, 559–569.

49. Petrosyan, V., Dobrolecki, L.E., LaPlante, E.L., Srinivasan, R.R., Bailey, M.H., Welm, A.L., Welm, B.E., Lewis, M.T., and Milosavljevic, A. (2022). Immunologically “cold” triple negative breast cancers engraft at a higher rate in patient derived xenografts. NPJ Breast Cancer 8, 104.

50. Perez-Riverol, Y., Bai, J., Bandla, C., García-Seisdedos, D., Hewapathirana, S., Kamatchinathan, S., Kundu, D.J., Prakash, A., Frericks-Zipper, A., Eisenacher, M., et al. (2022). The PRIDE database resources in 2022: a hub for mass spectrometry-based proteomics evidences. Nucleic Acids Res. 50, D543–D552.

51. Harvey, J.M., Clark, G.M., Osborne, C.K., and Allred, D.C. (1999). Estrogen receptor status by immunohistochemistry is superior to the ligand-binding assay for predicting response to adjuvant endocrine therapy in breast cancer. J. Clin. Oncol. 17, 1474–1481.

52. Saltzman, A.B., Leng, M., Bhatt, B., Singh, P., Chan, D.W., Dobrolecki, L., Chandrasekaran, H., Choi, J.M., Jain, A., Jung, S.Y., et al. (2018). gpGrouper: A Peptide Grouping Algorithm for Gene-Centric Inference and Quantitation of Bottom-Up Proteomics Data. Mol. Cell. Proteomics 17, 2270–2283.

53. Evrard, Y.A., Srivastava, A., Randjelovic, J., Doroshow, J.H., Dean, D.A., 2nd, Morris, J.S., Chuang, J.H., and NCI PDXNet Consortium (2020). Systematic Establishment of Robustness and Standards in Patient-Derived Xenograft Experiments and Analysis. Cancer Res. 80, 2286–2297.

54. Conway, T., Wazny, J., Bromage, A., Tymms, M., Sooraj, D., Williams, E.D., and Beresford-Smith, B. (2012). Xenome--a tool for classifying reads from xenograft samples. Bioinformatics 28, i172–8.

55. Li, H. (2013). Aligning sequence reads, clone sequences and assembly contigs with BWA-MEM. arXiv [q-bio.GN]. 10.48550/ARXIV.1303.3997.

56. DePristo, M.A., Banks, E., Poplin, R., Garimella, K.V., Maguire, J.R., Hartl, C., Philippakis, A.A., del Angel, G., Rivas, M.A., Hanna, M., et al. (2011). A framework for variation discovery and genotyping using next-generation DNA sequencing data. Nat. Genet. 43, 491–498.

57. McKenna, A., Hanna, M., Banks, E., Sivachenko, A., Cibulskis, K., Kernytsky, A., Garimella, K., Altshuler, D., Gabriel, S., Daly, M., et al. (2010). The Genome Analysis Toolkit: a MapReduce framework for analyzing next-generation DNA sequencing data. Genome Res. 20, 1297–1303.

58. Cingolani, P., Platts, A., Wang, L.L., Coon, M., Nguyen, T., Wang, L., Land, S.J., Lu, X., and Ruden, D.M. (2012). A program for annotating and predicting the effects of single nucleotide polymorphisms, SnpEff: SNPs in the genome of Drosophila melanogaster strain w1118; iso-2; iso-3. Fly *6*, 80–92.

59. Cingolani, P., Patel, V.M., Coon, M., Nguyen, T., Land, S.J., Ruden, D.M., and Lu, X. (2012). Using Drosophila melanogaster as a Model for Genotoxic Chemical Mutational Studies with a New Program, SnpSift. Front. Genet. 3, 35.

60. Karczewski, K.J., Francioli, L.C., Tiao, G., Cummings, B.B., Alföldi, J., Wang, Q., Collins, R.L., Laricchia, K.M., Ganna, A., Birnbaum, D.P., et al. (2020). The mutational constraint spectrum quantified from variation in 141,456 humans. Nature 581, 434–443.

61. Landrum, M.J., Lee, J.M., Benson, M., Brown, G.R., Chao, C., Chitipiralla, S., Gu, B., Hart, J., Hoffman, D., Jang, W., et al. (2018). ClinVar: improving access to variant interpretations and supporting evidence. Nucleic Acids Res. 46, D1062–D1067.

62. Kuilman, T., Velds, A., Kemper, K., Ranzani, M., Bombardelli, L., Hoogstraat, M., Nevedomskaya, E., Xu, G., de Ruiter, J., Lolkema, M.P., et al. (2015). CopywriteR: DNA copy number detection from off-target sequence data. Genome Biol. 16, 49.

63. Mermel, C.H., Schumacher, S.E., Hill, B., Meyerson, M.L., Beroukhim, R., and Getz, G. (2011). GISTIC2.0 facilitates sensitive and confident localization of the targets of focal somatic copy-number alteration in human cancers. Genome Biol. 12, R41.

64. Li, B., and Dewey, C.N. (2011). RSEM: accurate transcript quantification from RNA-Seq data with or without a reference genome. BMC Bioinformatics 12, 323.

65. Tyanova, S., Temu, T., and Cox, J. (2016). The MaxQuant computational platform for mass spectrometry-based shotgun proteomics. Nat. Protoc. 11, 2301–2319.

66. Wen, B., Jaehnig, E.J., and Zhang, B. (2022). OmicsEV: a tool for comprehensive quality evaluation of omics data tables. Bioinformatics 38, 5463–5465.

67. Jeffrey T. Leek and W. Evan Johnson and Hilary S. Parker and Elana J. Fertig and Andrew E. Jaffe and Yuqing Zhang and John D. Storey and Leonardo Collado Torres (2024). sva: Surrogate Variable Analysis. 10.18129/B9.bioc.sva.

68. Silver, D.P., Richardson, A.L., Eklund, A.C., Wang, Z.C., Szallasi, Z., Li, Q., Juul, N., Leong, C.-O., Calogrias, D., Buraimoh, A., et al. (2010). Efficacy of neoadjuvant Cisplatin in triple-negative breast cancer. J. Clin. Oncol. 28, 1145–1153.

69. Wolf, D.M., Yau, C., Wulfkuhle, J., Brown-Swigart, L., Gallagher, R.I., Lee, P.R.E., Zhu, Z., Magbanua, M.J., Sayaman, R., O’Grady, N., et al. (2022). Redefining breast cancer subtypes to guide treatment prioritization and maximize response: Predictive biomarkers across 10 cancer therapies. Cancer Cell 40, 609–623.e6.

70. Liao, Y., Wang, J., Jaehnig, E.J., Shi, Z., and Zhang, B. (2019). WebGestalt 2019: gene set analysis toolkit with revamped UIs and APIs. Nucleic Acids Res. 47, W199–W205.

71. Savage, S.R., Shi, Z., Liao, Y., and Zhang, B. (2019). Graph Algorithms for Condensing and Consolidating Gene Set Analysis Results. Mol. Cell. Proteomics 18, S141–S152.

72. Dowd, C. (2022). twosamples: Fast Permutation Based Two Sample Tests. Preprint.

