## Supplemental Figures S1-6 for "Patient-Derived Xenograft-Guided Prediction of Chemotherapy Response in Triple-Negative Breast Cancer"

### Figure S1: Lack of Enhanced Combination Response vs Best Single Agent

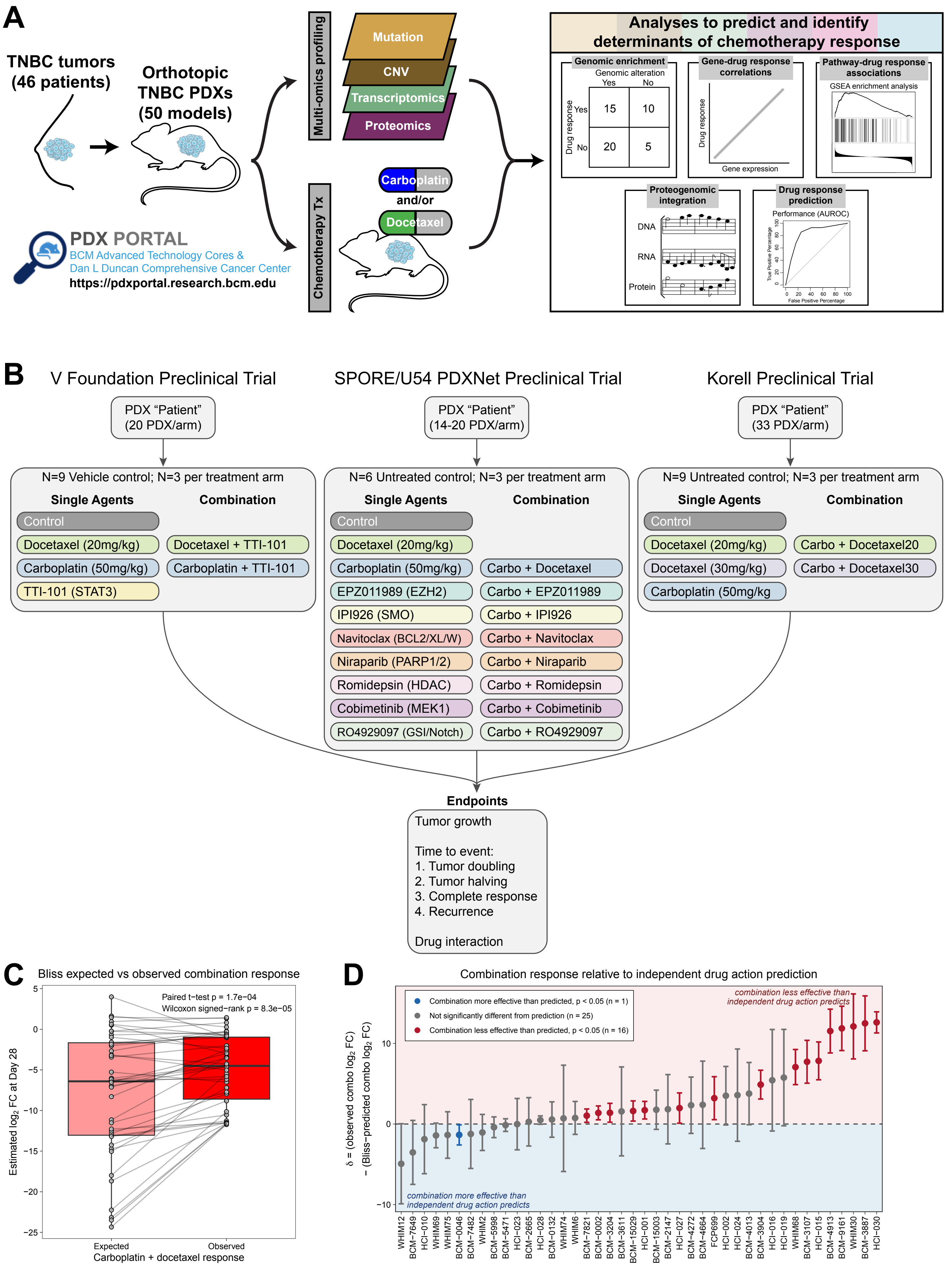

### Figure S2: Molecular Associates of Single Agent and Combination Treatment

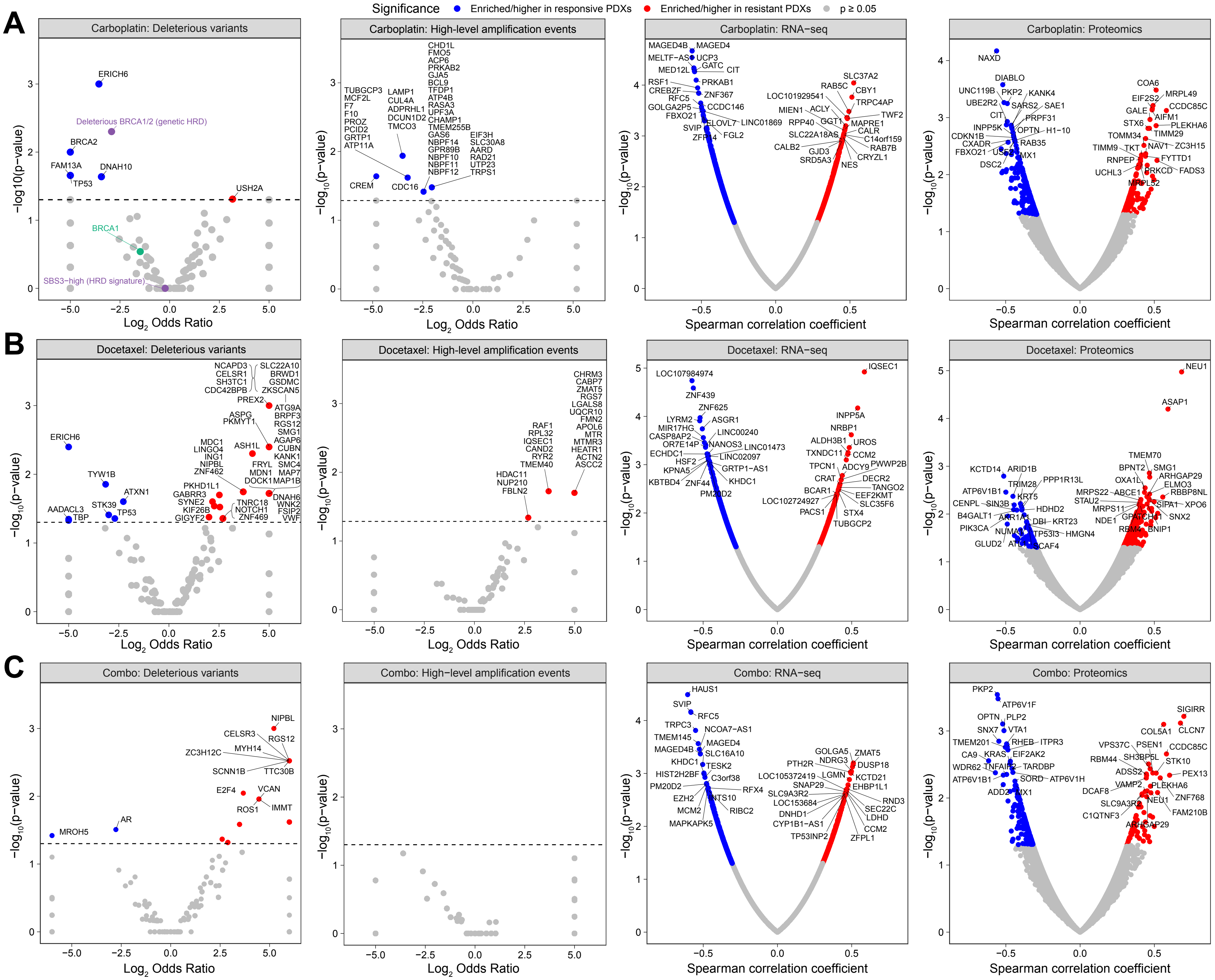

### Figure S3: Associations with Resistance to All Chemotherapy Arms

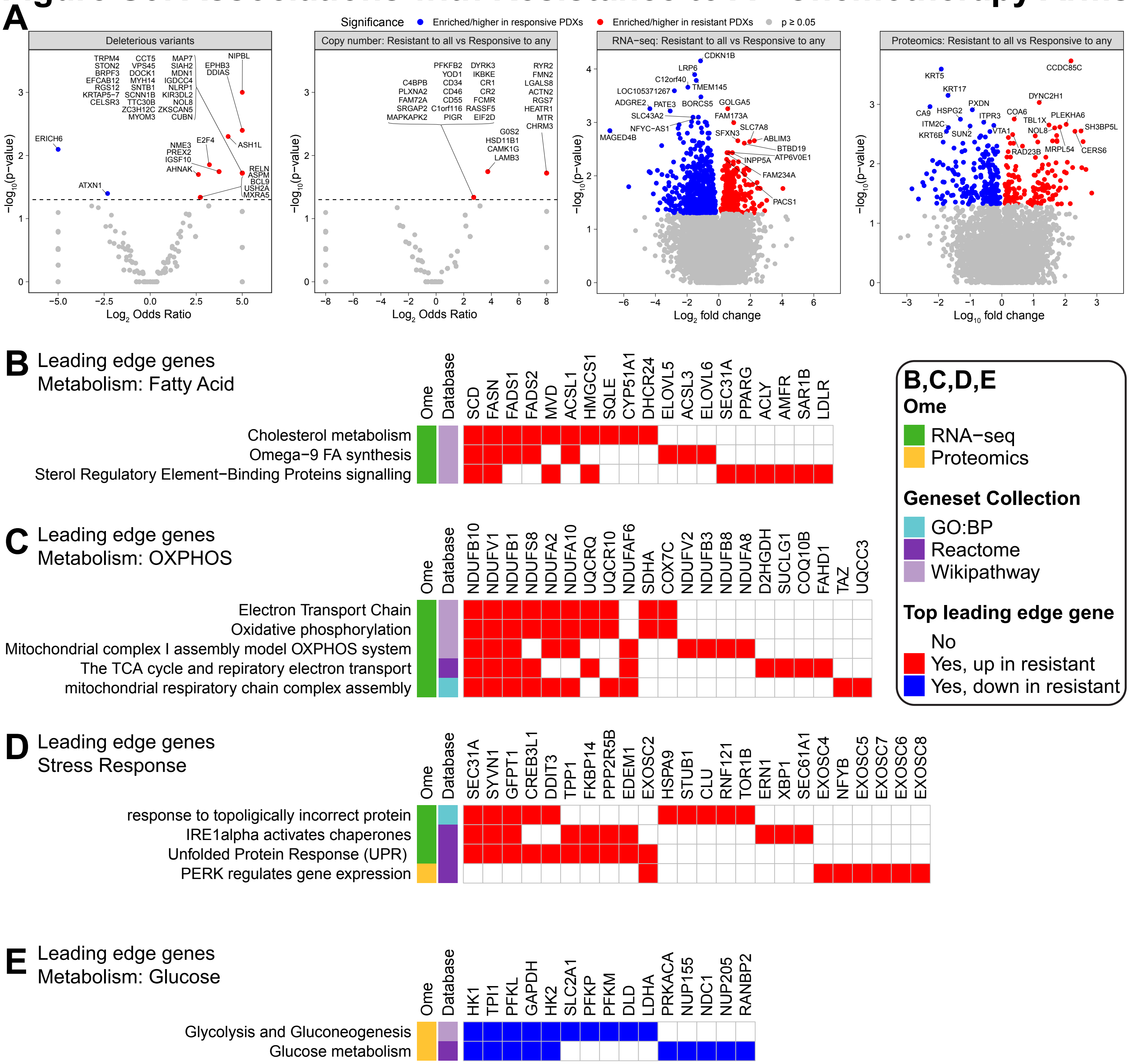

### Figure S4: Predicting Chemotherapy Response

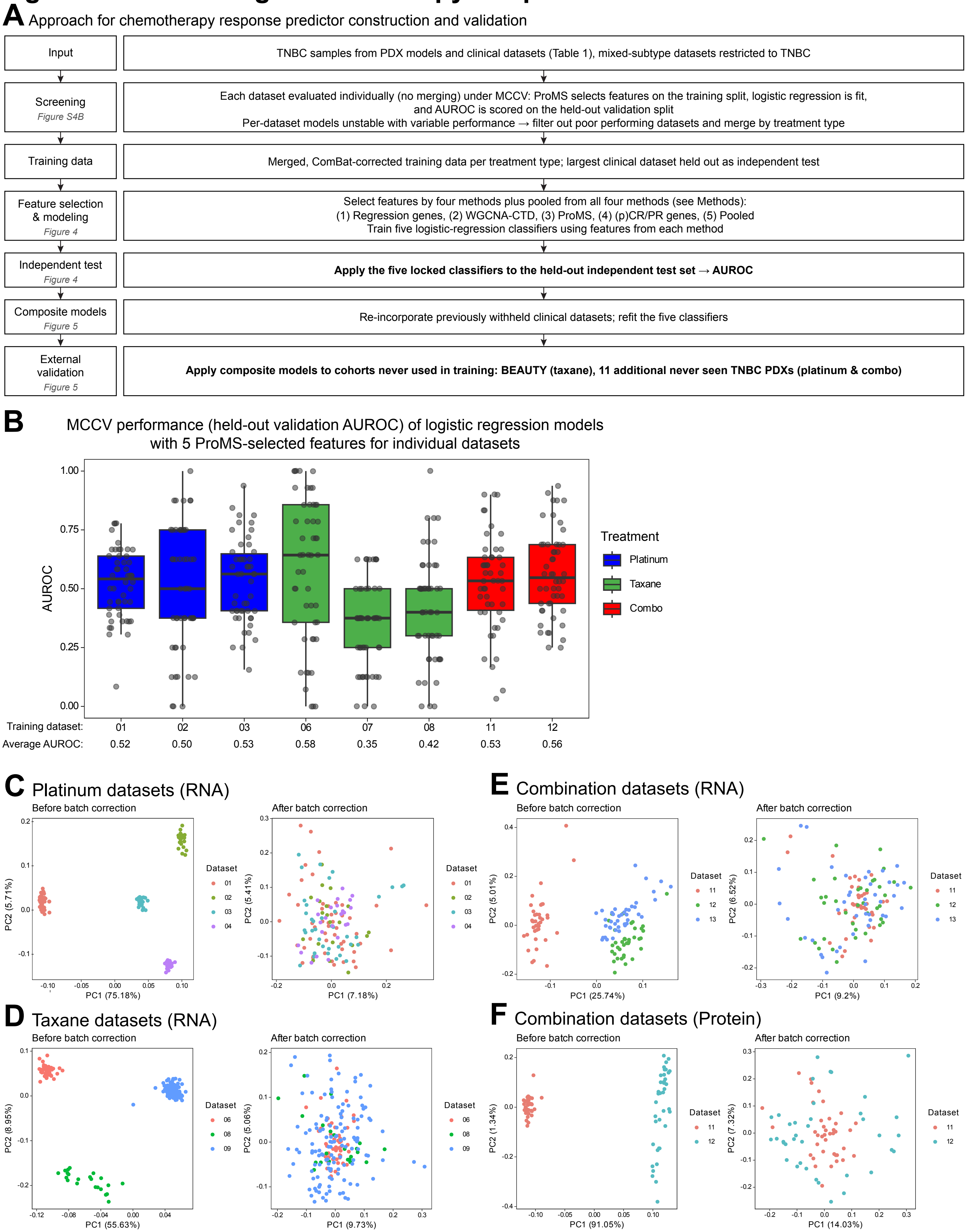

### Figure S5: ProMS for Single- and Multi-omic Predictors of Chemoresponse

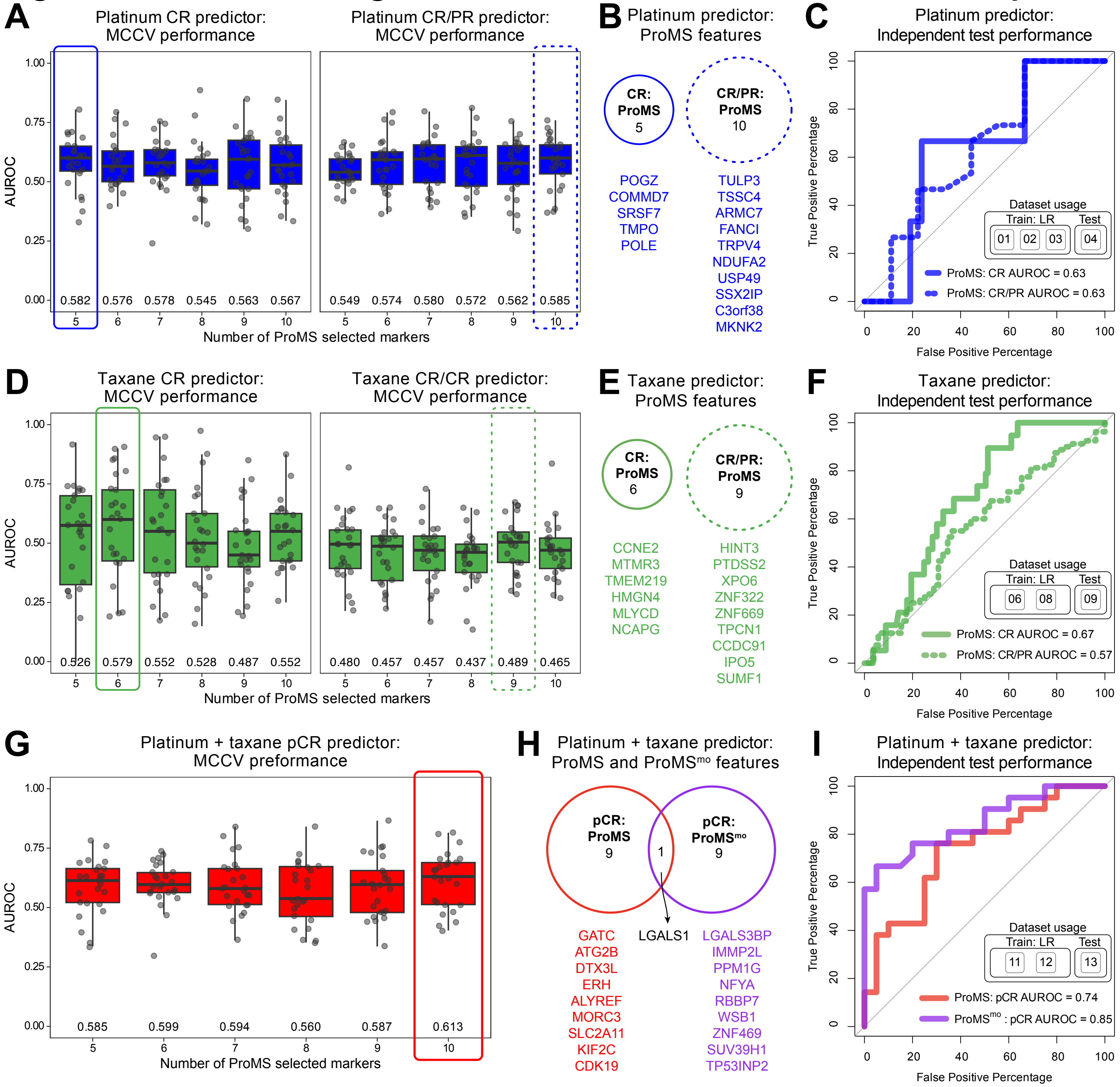

### Figure S6: Association of Predictive Features with Gene Function

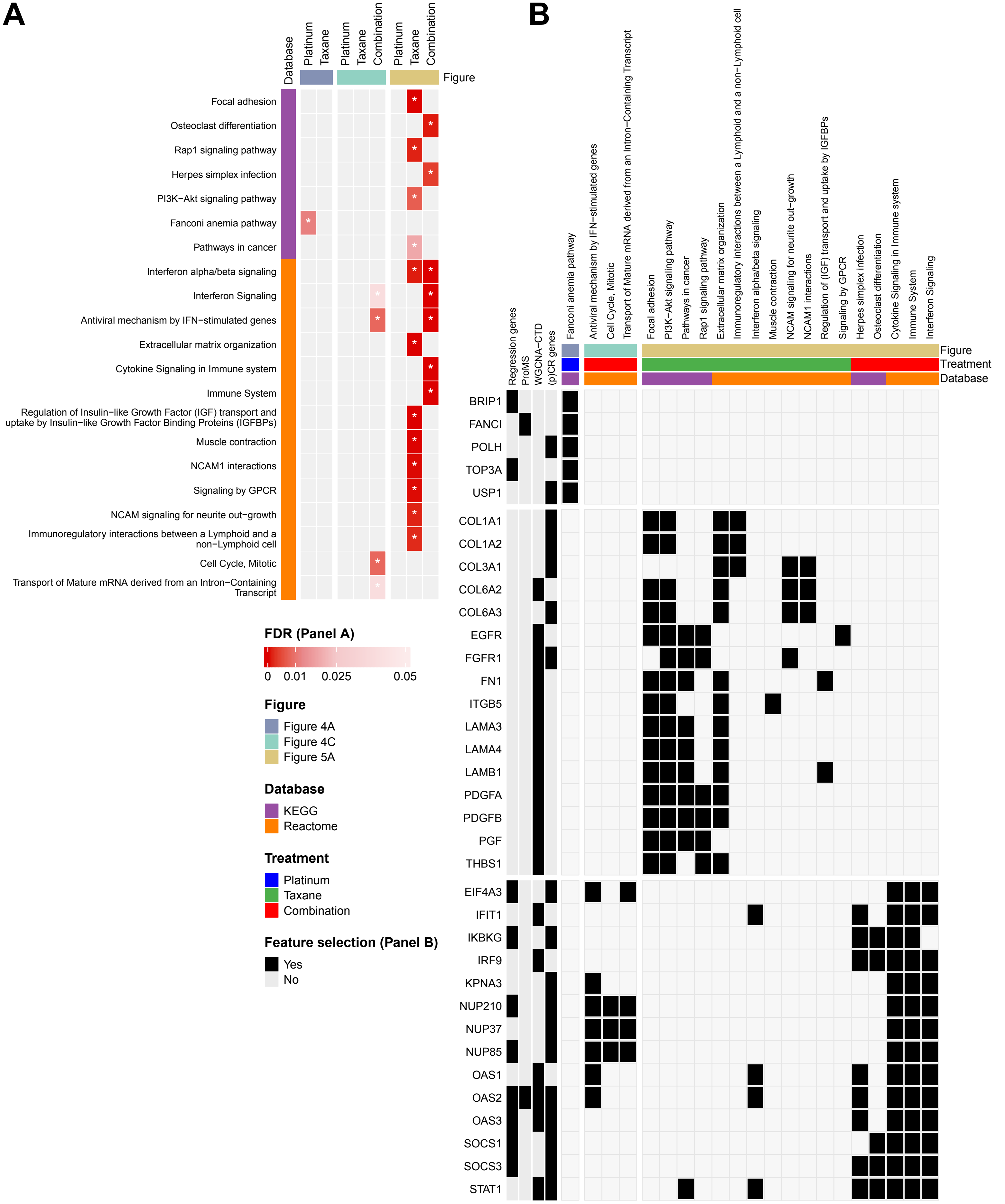
